# Three-Dimensional Piezoelectric Fibrous Constructs for Muscle Regeneration

**DOI:** 10.1101/2025.06.19.660455

**Authors:** Xenofon Karagiorgis, Gerasimos Balatsoukas, Mahdieh Shojaei Baghini, Merna Maung, Jiaqi Zong, Shruti Niar, Charchit Kumar, Aleixandre Rodrigo-Navarro, Daniel M. Mulvihill, Peter J. Skabara, Carlos Garcia Nuñez, Oana Dobre

## Abstract

Poly(3-hydroxybutyrate-co-3-hydroxyvalerate) (PHBV), a biocompatible and biodegradable polymer with inherent piezoelectricity, holds promise for developing biomimetic scaffolds for tissue regeneration. Its ability to generate electrical charges in response to mechanical stimulation, mimicking the electromechanical environment in native tissues like muscle, makes it attractive. This study focuses on harnessing PHBV properties to engineer electrospun fibrous scaffolds for muscle regeneration. We investigated processing parameters, such as solvent combinations, polymer concentrations, and tip-to-collector distances on fiber morphology, porosity, and piezoelectric properties. We observed a 25-fold increase in the quasi-static piezoelectric coefficient (d33 from 0.117 to 2.9 pC/N) for scaffolds with controlled porosity. This piezoelectric activity was corroborated by dynamic Vector Network Analyzer (VNA) measurements, which identified an electromechanical coupling factor (kt) of 13.86%. This enhanced electromechanical response highlights their potential for stimulating cellular responses for tissue regeneration. The mechanical properties of these scaffolds demonstrated their ability to undergo strain up to 40%, within the optimal range for muscle tissue engineering and compatibility with muscle functionality. Cytocompatibility of PHBV scaffolds was confirmed, and preliminary studies suggest their potential to support myogenic differentiation. This research highlights PHBV-based biomimetic scaffolds potential for muscle regeneration and sets groundwork for developing advanced therapies addressing muscle injury and disease.

## 1. INTRODUCTION

Skeletal muscle tissue comprises approximately 40% of total body mass and plays crucial roles in locomotion, posture maintenance, and protection of internal organs. Severe muscle injuries, particularly those resulting in volumetric muscle loss (VML) where more than 20% of muscle volume is compromised, pose significant clinical challenges due to the limited regenerative capacity of adult skeletal muscle,^1^ Following such substantial tissue loss, the body’s endogenous repair mechanisms prove insufficient, leading to persistent inflammation, extensive scarring, and compromised functional recovery.^2,3^ Current clinical interventions for VML predominantly rely on autologous muscle grafts, which have demonstrated positive outcomes in supporting muscle regeneration.^4^ However, these approaches present several limitations, including donor site morbidity and suboptimal tissue integration due to random fiber orientation resulting from the necessary mincing process.^5^ The compromised structural organization of these grafts often leads to extensive scar tissue formation, thereby limiting functional recovery. Recent advances in tissue engineering have led to the development of three-dimensional (3D) fibrous piezoelectric scaffolds (FPS) as potential alternatives to traditional grafts.^6,7^ These biomimetic constructs combine structural support with electromechanical stimulation, closely replicating the native muscle environment.^8,9^ These 3D scaffolds based on synthetic polymers, e.g., polyvinylidene fluoride (PVDF) or poly(3-hydroxybutyrate-co-3-hydroxyvalerate) (PHBV), the latter proposed in this work, naturally exhibiting piezoelectricity due to their molecular structure lacking of center of symmetry (**Figure 1a and b**) enabling the generation of electrical signals in response to mechanical deformation. This unique feature mimics the natural electromechanical coupling essential for muscle function and regeneration (**Figure 1(c)**).^10,11^ Another advantageous characteristic of the FPS is the mimicry of the natural extracellular matrix (ECM) microarchitecture of various tissues while engineered to exhibit excellent mechanical properties.^12^ The higher surface area of the fibers plays a vital role in the exchange of nutrients and oxygen which is particularly beneficial for applications in load-bearing tissues, such as muscles. ^13,14^

**Figure 1.**
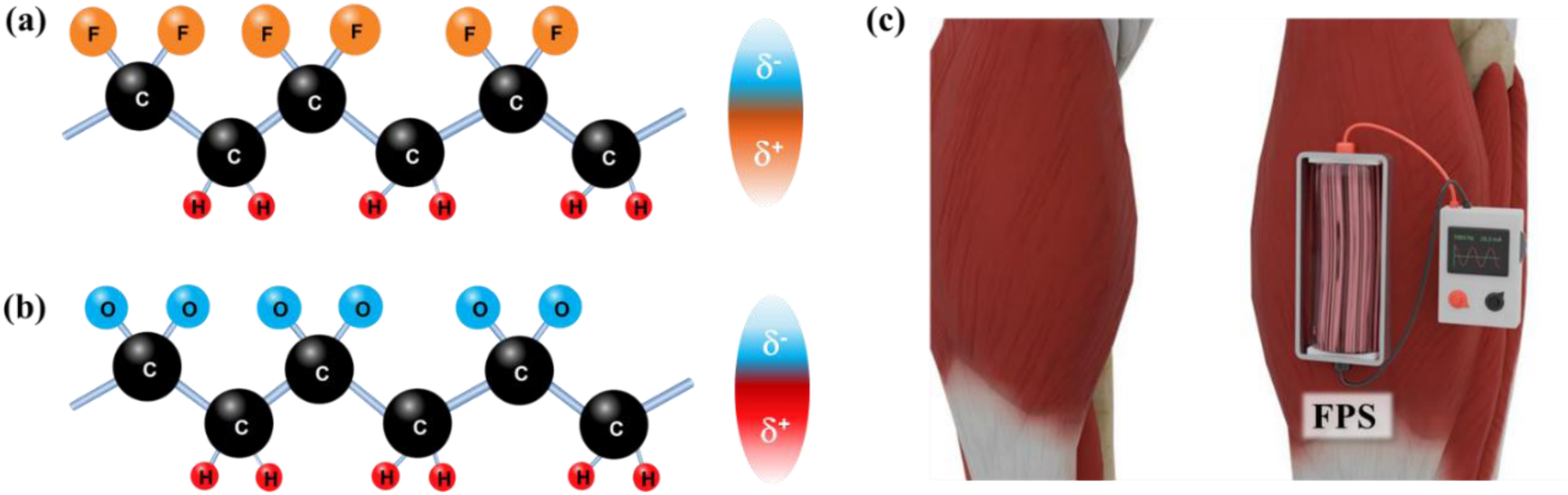
Schematic of the molecular structure of two piezoelectric synthetic polymers, including (a) PVDF and (b) PHBV showing the electric dipole form upon mechanical deformation of the molecule due to non-central symmetry. (c) Drawing of FPS used for muscle regeneration that can be local stimulation applied by the vibrating fibers.

Among various fabrication techniques for three-dimensional fibrous constructs (including self-assembly^15^ and phase separation^16^), electrospinning^17,18^ has emerged as the preferred method due to its versatility and cost-effectiveness.^19^ This technique enables precise control over fiber dimensions and morphology within the micro- and nano-meter range, while facilitating functionalization with bioactive molecules.^20^ The resulting scaffolds exhibit high surface area and interconnected porosity, crucial features for cellular infiltration and nutrient exchange in muscle tissue engineering applications. PHBV, a member of the polyhydroxyalkanoates (PHAs) family, has attracted significant attention as a promising material for tissue regeneration applications.^21,22^ This natural aliphatic thermoplastic polyester combines biodegradability and biocompatibility with inherent piezoelectric properties, and comprises vertical (*d*_33_) and lateral (*d*_31_) piezoelectric coefficients of 2-5 pC/N and 0.5-2 pC/N, respectively^23^, which are similar to the piezoelectric coefficient of the natural bone. PHBV’s *in vivo* degradation produces metabolically compatible byproducts, while its mechanical properties and piezoelectric crystalline phase support both structural and electrical aspects of muscle regeneration.^24^

Despite these advantages, the optimization of PHBV-based three-dimensional piezoelectric fibrous scaffolds remains challenging. Existing approaches often necessitate complex modifications, including the incorporation of additional polymers^25,26^ or nanostructures^27,28^, and post-processing treatments.^29^ These requirements not only increase production complexity but also raise concerns regarding scalability and clinical translation. The present study addresses these challenges by developing a streamlined approach for fabricating PHBV-based three-dimensional piezoelectric fibrous constructs specifically designed for muscle regeneration. We investigate the influence of processing parameters on fiber morphology, piezoelectric properties, and biological performance, aiming to establish an optimized protocol for producing scaffolds that effectively support muscle tissue regeneration while maintaining simplicity and reproducibility in fabrication.

## 2. MATERIALS AND METHODS

### 2.1 Materials

PHBV with an 8% molar ratio of hydroxyvalerate (HV) was purchased from Biopol and Sigma Aldrich. Chloroform, dichloromethane (DCM), and dimethylformamide (DMF) were purchased from Sigma-Aldrich. All chemical reagents were used as received, without further purification.

### 2.2 PHBV Solution preparation

PHBV granules were added to different solvents, as shown in **Table S1**, and stirred overnight at 60 ℃/200 rpm until a homogenous solution was obtained.

### 2.3 Electrospinning

All solutions suitable for electrospinning were loaded into 10 mL syringes within the electrospinning setup (TL-PRO, TONGLI TL, China), as shown in **Figure 2a**. The experiments were conducted at a consistent 40% ± 3% relative humidity and a temperature of 25°C. A grooved collector, rotating at 1000 rpm and connected to an electrical voltage of 0.3 kV, was used to collect the align electrospun fibers (*see* Video S1 in the supplementary information). The presence of grooves on the collector surface generated a non-uniform electric field distribution. Electric field lines were concentrated at the groove edges, establishing preferential deposition sites that guided fiber orientation. This mechanism encouraged fibers to bridge across the groove gaps, stretching between the raised edges (ridges) of adjacent grooves, thereby naturally aligning them perpendicularly to the groove direction.

**Figure 2.**
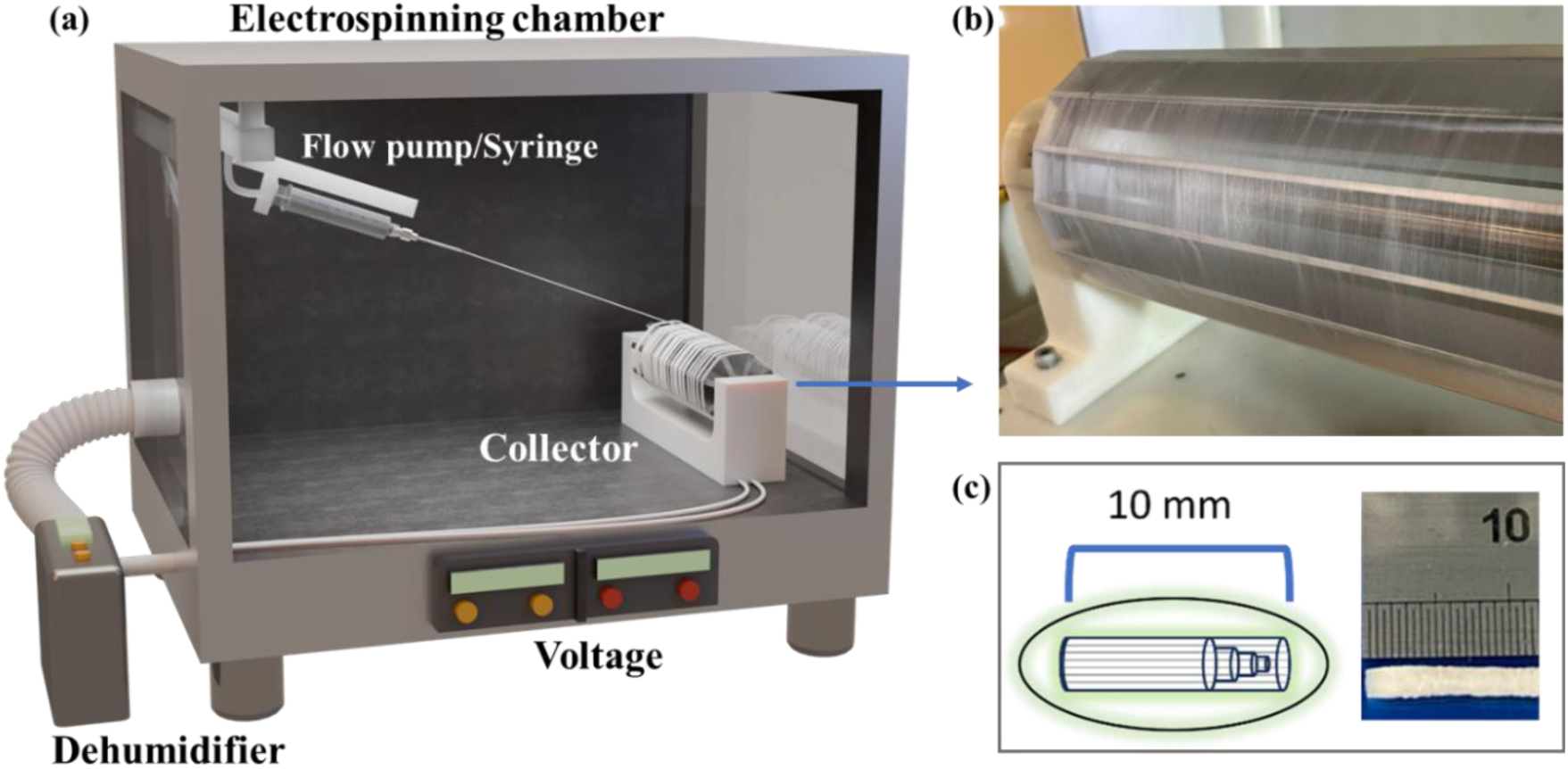
(a) Schematic illustration of the electrospinning setup. (b) PHBV fibers deposited on the grooved collector and (c) PHBV rolled cylindrical scaffold schematic drawing and a picture (5X).

The solutions were supplied at a rate of 1 mL/h through a 20G stainless steel needle, positioned 15 cm away from the collector. The applied voltage varied for each solution, as detailed in **Table S2**, representing the minimum electrospinning voltages (MEVs) needed to form visible Taylor cones. To achieve uniform, align fiber distribution, the syringe system was programmed to scan at a speed of 40 mm/s, ensuring even material deposition across the collector. The deposition time was maintained at 8 hours to ensure enough quantity of electrospun fibers, as demonstrated in **Figure 2b** and then rolled as shown in **Figure 2c**.

### 2.4 Fibers characterisation

The morphology of the fibers was characterized in a JEOL IT 100 scanning electron microscope (SEM), with an accelerating voltage of 10 kV and imaged using InTouchScope software. A sheet of aluminum foil was placed on the collector before producing fibers to be able to mount the samples on the SEM holder Prior to the characterisation, the deposited fibers on aluminum foil were sputter coated with 15 nm of gold. The FEI Nova NanoSEM 450 with a voltage of 5kV was used to measure the thickness of the fibers. The Instron 3000 linear-torsion all-electric dynamic test instrument was used to assess the mechanical properties of the PHBV fibers. The chemical groups of PHBV were identified using an Agilent Cary 630 Fourier-Transform Infrared (FTIR) spectrometer. The spectrometer spectra range is 650 to 4000 cm^-1^ with a resolution of 2 cm^-1^.

#### 2.4.1. Berlincourt method

The piezoelectric characterization of PHBV fibres was carried out by using an optimized version of a Berlincourt (BC) d33 meter (PM-300 from Piezotest Ltd.). For all measurements, and following previous research results, the static force (*F*_s_) was fixed to 1 N, the dynamic force (*F*_d_) was set to 0.25 N, and the measuring frequency (*f*) was set at 110 Hz. To ensure the reproducibility and reliability of the measurements, the samples were measured for 10min, then removed from excitation, and remeasured. The cycle was repeated up to five times, and the piezoelectric coefficient was averaged to obtain the displayed result.

#### 2.4.2. Piezoelectric coefficient measurements using Vector Network Analyzer (VNA) method

The high frequency RF properties of the electrospun nanofibers were characterized in a high-precision Vector Network Analyzer (Keysight N5245B) following short-open-load calibration. The interdigitated electrodes (IDE) structures were soldered to an unterminated-SMA (50 Ω) cable prior to connection. The intermediate frequency (IF) bandwidth was kept constant at 1 kHz with an input power of 0 dBm. The impedance and phase were obtained from the recorded reflection parameters. The core resonance region was identified between 1.74 and 1.8 GHz, during which the PHBV nanofibers exhibit a phase transition from capacitive to inductive behavior near resonance, returning to capacitive behavior around anti-resonance. The measurements were carried out in conjunction with a blank IDE control sample to rule out the peaks associated with IDE resonance.

##### Patterning of interdigitated electrodes using VNA method

Copper tape (Advanced Tape) with a thickness of 65±3μm was patterned using a precision blade-cutting method (Cameo 4 from Silhouette America) to create IDE structures. This device allows precise electrode shape design and automated cutting, ensuring high accuracy and reproducibility. The geometry of the electrodes was carefully optimized to maximize the electric field interaction with the PHBV fibers, enhancing their electrical response. The final electrode design, as fabricated using this method, is shown in **Figure 3a**, illustrating the well-defined interdigitated structure used for impedance characterization. The IDE dimensional parameters were as follows: finger length (*L*) of 34.1 mm, finger width (*W*) of 2 mm, inter-electrode gap (*G*,*G*_e_) of 2 mm, number of fingers of 14, and an electrode pitch of 8 mm.

**Figure 3.**
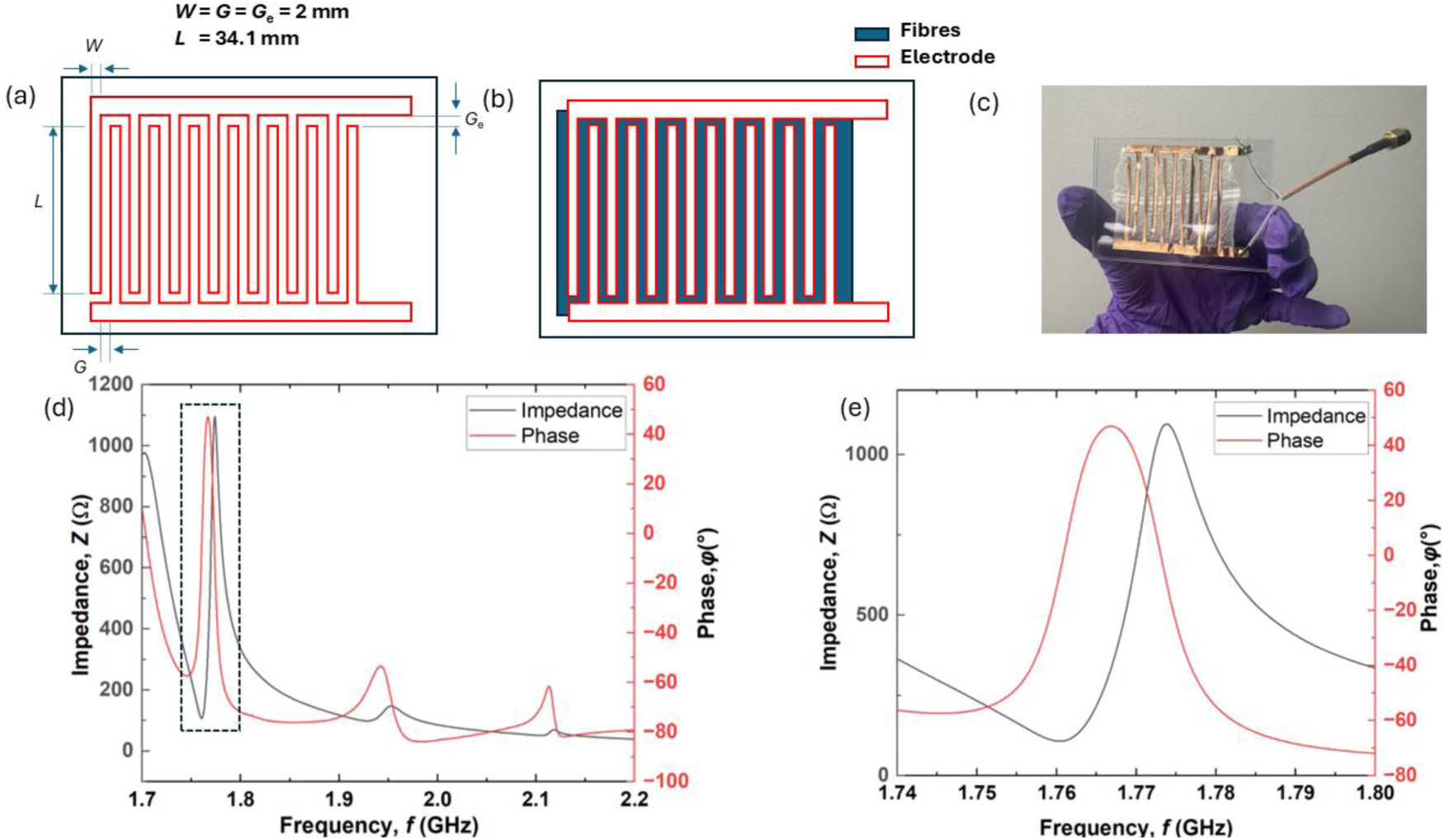
Schematic diagrams of the (a) IDE design and its dimensional parameters, and (b) the IDE with the fibers. (c) Photograph of the final device with a SMA connector. (d) Impedance and phase parameters vs frequency. (e) zoom-in of the first resonant mode.

##### Transfer of PHBV fibers on IDE

Firstly, the PHBV were gently transferred onto a glass slide. Then, the prepared IDE structure was mechanically pressed against the surface of the fibers, and thanks to the conductive glue of the tape, they form a stable electric contact (**Figure 3b**). Special care was taken during this process to align the electrodes as parallel as possible and ensure proper contact with the fiber surface. **Figure 3c** shows a photograph of the resulting device with a SMA connector that was attached to the Cu electrodes allowing the testing of piezoelectric properties in a VNA.

##### Electromechanical coupling factor (**𝒌_𝒕_^𝟐^**) and piezoelectric coefficient (**𝒅_𝟑𝟑_**) calculation

The electromechanical coupling coefficient (𝑘_𝑡_^2^) quantifies the efficiency of energy conversion between electrical and mechanical domains within the PHBV fiber structure. It was calculated using the measured resonance frequencies of a *thickness extensional mode* (IEEE Standards ANSI/IEEE 176 -1987) following **Equation 1**:

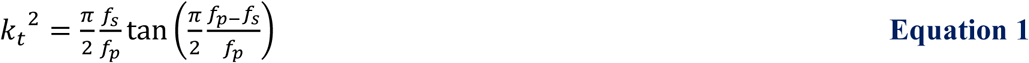

A higher 𝑘_𝑡_^2^ value indicates a more effective piezoelectric response, which is critical for optimizing the performance of PHBV fibers in bioelectronic applications.

### 2.5 FEM Analysis of Pore Size Effects on Piezoelectric Response in PHBV Nanofibers

The displacement distribution of the PHBV fibers with different amount of pore volume was simulated using COMSOL Multiphysics version 6.2.^30^ The fiber was modelled as a 3D cylindrical geometry with a length of 70 µm, a radius of 790 nm, and a total volume of 1.37 × 10^11^ nm^3^. The mesh is free tetrahedral with a minimum element size of 15 nm. Three cases were considered: a non-porous fiber (0 pores) and porous fibers containing either 100 or 1500 pores within the same volume. These simulations aimed to provide insights into how different pore volumes influence displacement distribution. Boundary conditions were applied during the test to analyze the stress distribution, with the bottom surface fixed as a constraint. The values of *d*_33_ and Young’s modulus (*E*) were obtained through experimental characterisation. The Poisson’s ratio (*v*) and density (*ρ*) were derived from a previous study^31^ and are listed in **Table S3.**

### 2.6 Tensile testing

Uniaxial tensile testing was performed using an electrodynamic tensile testing machine (Model E3000, Instron UK) to characterize the mechanical properties of the electrospun materials. Two distinct sample configurations were investigated: flat electrospun fiber mats (thin sheets) and 3D cylindrical scaffolds formed by rolling the electrospun fibers.

Flat fiber mats were attached using soft adhesive tape and tightening screws and attached to the clamps. Prior to each test, the sample’s alignment within the grips was meticulously checked to ensure uniform loading and proper orientation along the central axis.

Measurements were performed under displacement control at a crosshead speed of 10 mm/min. Force and displacement data were continuously recorded throughout the tests, enabling the derivation of stress-strain curves.

The 3D cylinders rolled fibers samples were clamped using a pair of custom-designed clamps to ensure the even distribution of tensile forces during testing. To adjust for any difference between the clamping locations of the samples, the length of the sample was re-measured once the sample was clamped. Both clamps were attached to the rest of the assembly via magnets, with one clamp being directly attached to the load cell and the other to a custom-designed ‘arm’ that allowed for free motion in the y and z planes for better alignment of the clamps. The platform was then modified to be able to stand on its side, and a 3D-printed bath was custom designed to encapsulate the two clamps. This allowed the clamps to be submerged in liquid while conducting tests, which was useful to evaluate the mechanical properties of the fibers in wet conditions. The liquid used in the water bath during the wet testing was the same solution used for media in the culturing of C2C12 cells, which consisted of 80% DMEM (Dulbecco’s Modified Eagle’s Medium (DMEM, Gibco), 20% FBS (fetal bovine serum) and 1% P/S (Penicillin/Streptomycin). The entire setup for the tensile tests can be seen in **Figure S1**.

### 2.7 Cell viability assessment

PHBV scaffolds were subjected to oxygen plasma treatment at 80 W for 3 minutes under vacuum conditions to increase the hydrophilicity of the scaffold. This was followed by UV-sterilization of each side of the scaffold for 30 mins each. 20 µl of 20 µg/ml Laminin 211(LM211) (BioLamina) was then absorbed onto three of the scaffolds (treated) for 1 h, while the other three scaffolds were left uncoated. Laminins (LMs) are important structural proteins of the extracellular matrix (ECM).^32^ The abundance of every LM isoform is tissue-dependent, suggesting that LM has tissue-specific roles.^33^ Laminin 211 is found in the basement membrane surrounding the sarcolemma has a role in muscle function. Then PHBV scaffolds were then placed in a 48-well non-treated tissue culture plate and seeded with 20000 C2C12 cells and incubated at 37°C with 5% CO_2_ for three time points: Day 1, Day 3, and Day 7.

Once it reached each time point, the media was aspirated from the cells, and the cells were incubated with 150 µL of a dye mixture from the LIVE/DEAD™ Viability/Cytotoxicity Kit (Invitrogen™) for 20 minutes in the incubator. The dye mixture was prepared by diluting Calcein AM (1/500) and Ethidium homodimer (1/2000) in PBS. After incubation, the scaffolds were washed with PBS, and 200 µL of DPBS was added to each well to prevent the scaffolds from drying out before imaging in a Zeiss LSM-900 confocal microscope.

### 2.8 Cell-matrix interaction

PHBV scaffolds were prepared as in Section 2.7 for oxygen plasma treatment, UV-sterilisation, laminin coating and C2C12 seeding. The cells were then incubated for 3 hours in the incubator with serum-free media. C2C12 cells plated on the fibers were fixed in 4% formaldehyde for 20 minutes, followed by a wash with DPBS, and permeabilized at room temperature with 0.1% Triton/DPBS for 15 minutes. The samples were then blocked to prevent non-specific background binding at room temperature with 1% BSA/DPBS for 30 minutes.

Next, the samples were incubated with a mouse monoclonal anti-vinculin primary antibody (Sigma, 1:400) for 1 hour at room temperature. Subsequently, the samples were washed with 0.5% Tween/DPBS for 5 minutes, 5 times, and stained with secondary Cy3 rabbit anti-mouse antibody (Jackson ImmunoResearch, 1:200), together with phalloidin (Sigma,1:100) and DAPI (Sigma, 1:500), for 1 hour. This was followed by five 5-minute washes on a rotating plate. Finally, DPBS was added to the samples, and they were stored at 4°C for microscopy.

### 2.9 Myogenic differentiation

As described in Section 2.8, PHBV scaffolds were treated with oxygen plasma, UV sterilise, Laminin absorption, and placed in 48 well, non-treated tissue culture plates. 5000 of C2C12 cells were seeded on each sample and incubated to grow until it reached 100% confluency. Upon reaching the confluency, the growth media was switch with an induction media consisting of components such as (DMEM+ 1% Pen/Strep + 0.1% FBS + 1% Insulin Transferrin Selenium). The cells were incubated with the differentiation media for nine days before conducting q-PCR or immunostaining.

### 2.10 qPCR - Gene Expression

C2C12 cells that were growing on the PHBV scaffolds were detached using 1x trypsin-EDTA in a 2ml Eppendorf. Total RNA was then extracted from the detached cells using RNeasy mini kit (Qiagen) and quantified with a Nanodrop 1000 (ThermoScientific). Following this, extracted RNA from all samples were normalised to the sample consisting of lowest RNA concentration. Subsequently, following the manufacturer’s instruction, genomic DNA was wiped out and RNA was reverse transcribed to cDNA using a QuantiTect Reverse Transcription Kit (Qiagen). qRT-PCR was conducted using a 7500 real-time PCR system (Applied Biosystems) and SYBR Green master mix. PCR cycle reaction for 40 cycles were as follows: 50°C for 2 min followed by 95° C for 10 min and then amplified at 95° C for 15 s and 60° C for 1 min. The sequences for the Human-specific primers employ for qPCR amplification were as follows: MyoD (Gene ID: 17927; Fw: 5′-GCACTACAGTGGCGACTCAGAT-3′, Rev: 5′-TAGTAGGCGGTGTCGTAGCCAT-3′), Myogenin (Gene ID: 17928; Fw: 5′CCATCCAGTACATTGAGCGCCT-3′, Rev: 5′-CTGTGGGAGTTGCATTCACTGG-3′), GAPDH (Gene ID: 14433; Fw: 5′-CATCACTGCCACCCAGAAGACTG-3′, Rev: 5′-ATGCCAGTGAGCTTCCCGTTCAG-3′) was used as a housekeeping gene.

## 3. RESULTS and DISCUSSION

### 3.1 Characterisation of PHBV fibers morphology

The optimal combination and concentration of the solution to produce PHBV fibers were determined by conducting morphological characterisation. The average fiber and bead diameters were measured using ImageJ software by analyzing 15 individual fibers for each PHBV solution. **Figure 4** presents the morphology of PHBV fibers using Biopol-supplied polymer. In **Figure S2a, b, and e**, it is evident that only the combination of chloroform with DCM in various v/v ratios produced PHBV solutions with visible fibers. In the case of B5_CD1 fibers, fibers with beads were observed, with an average diameter of 0.91 μm ± 0.31 μm. B5_CD2 exhibited fibers with slightly higher diameter of 0.97 μm ± 0.21 μm, also with some beads. However, for B5_DF and B5_CF (**Figure S2c and 3d)**, only beads were detected without any fibers. When the concentration of the solution was increased from 5 wt% to 8 wt% (B8_CD2), bead-free fibers were formed with an average diameter of 1.27 ± 0.19 μm. This outcome may be attributed to the viscosity of the solution, as it is proved that can regulate the flow of the solution and thus the formation and morphology of the fibers.^34^

**Figure 4.**
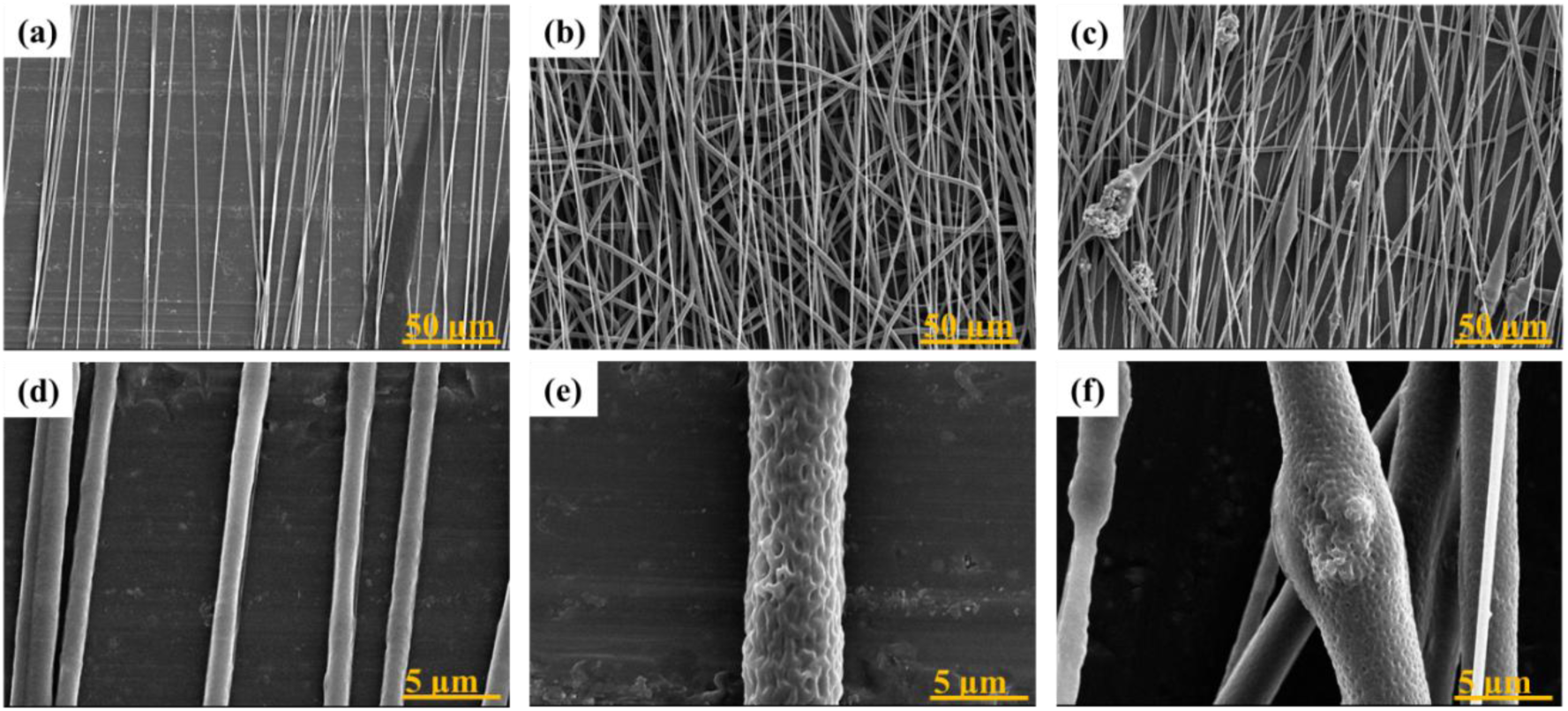
SEM images of the B8_CD2 fibers at (a) 15 cm, (b) 20 cm, (c) 30 cm tip-to-collector distances and higher magnification of B8_CD2 fibers at (d) 15 cm, (e) 20 cm and (f) 30 cm.

To further investigate the effect of tip-to-collector distance, the distance was increased from 15 cm (**Figure 4a**) to 20 and 30 cm, as shown in **Figure 4b and 4c**, respectively. The results indicated that increasing the tip-to-collector distance led to the formation of larger electrospun fibers with porous structures, as shown in **Figure 4e** and **4f**. The average fiber diameters measured at distances of 15 cm, 20 cm, and 30 cm were 1.27 ± 0.19 μm, 1.58 ± 0.44 μm, and 2.09 ± 0.66 μm, respectively. Moreover, the fibers obtained at 20 cm and 30 cm exhibited porous structures, with average pore sizes of 0.29 ± 0.15 μm and 0.15 ± 0.05 μm, respectively. In the case of the fibers obtained at 30 cm, the formation of some beads with an average diameter of 8.46 ± 4.72 μm was also noticed. This may be due to the weaker electric field strength at this distance, which affects the stability of the electrospinning process.^35^ The formation of the beads embedded on the fibers may lead to weaker mechanical properties which in turn will make the scaffold less suitable for mimicking the natural ECM of muscle regeneration. The change in fiber morphology can be primarily attributed to reduction of the electrostatic force applying on the polymer jet. As the distance increases, the strength of the electrostatic field decreases which leads to less stretching of the fibers before they solidify. As a result, the obtained fibers at 20 cm and 30 cm have larger diameters. This effect is closely related to the voltage-to-distance (V/d) ratio, where a lower field strength weakens the pulling force on the jet. In this case, the applied voltage was kept constant at 15 kV, but this was not enough to compensate for the longer distance between the needle and the collector. As a result, the electrostatic force pulling the fibers became weaker which caused less stretching of the fibers during electrospinning.^36^ This reduced elongation led to an increase in fiber diameter. More details on the diameters of the produced fibers can be seen at **Table S2** and **Figure S3**. SEM cross-sectional images were used to analyze the thickness of the film based on smooth (15cm) and porous (20cm) fibers. It was found that the smooth fibers-based layer has 74.07 μm thickness, while the layer based on porous fibers have a thickness of 52.75 μm, as shown in **Figure S4**.

Regarding the PHBV obtained from Sigma Aldrich, the SEM images of the fibers are shown in **Figure S5**. In all cases using different solvents, fibers formed beads while in other cases only beads were formed. The most fibers with the less beads were observed when a solvent mixture of DCM and DMF (9:1 v/v) was used (SA5_DF), with an average diameter of 0.44 μm ± 0.16 μm for the fibers and 2.39 µm ± 1.25 μm for the beads. To address this issue, attempts were made to improve the PHBV solution in these solvents by increasing the polymer concentration to 8 wt% and 10 wt% to achieve a more suitable viscosity. In **Figure S5e and S5f**, it can be observed that as the PHBV concentration increased, the average fiber diameter decreased to 0.498 ± 0.2 μm and 0.38 ± 0.18 μm, respectively, but more beads were also produced. The increase in bead formation is possibly attributed to the instability of the Taylor cone. During the formation of fibers using the SA8_DF and SA10_DF solutions, it became challenging to maintain a continuous and uniform flow of the solution from the spinneret. This resulted in the production of thinner fibers with an increased number of beads. It is worth noting that the optimisation of fibers from this supplier was halted because when using BioTek-supplied PHBV, smooth and uniform fibers were obtained without issues related to bead formation.

### 3.2 FTIR analysis of electrospun PHBV fibers

The chemical groups in electrospun PHBV fibers obtained at different tip-to-collector distances, were identified through FTIR characterization. The B8_CD2 fibers were specifically selected for analysis as they exhibited the most uniform morphology, with minimal or no bead formation. **Figure 5** illustrates that all samples showed similar curves. More specifically, the peak at 1723 cm^-1^ indicated the ester carbonyl (C=O) of the PHBV polymer, and the 1380 cm^-1^ peak was associated with the symmetrical wagging of aliphatic methyl (CH_3_) in the PHBV polymer. The symmetric stretching vibration peaks of C-O in PHBV were observed at 1453 cm^-1^ and 1054 cm^-1^. It was also observed that all typical PHBV peaks were present in all forms, and there were no observable shifts, suggesting that there is no alteration of the chemical groups of the material under different electrospinning parameters.^37^

**Figure 5.**
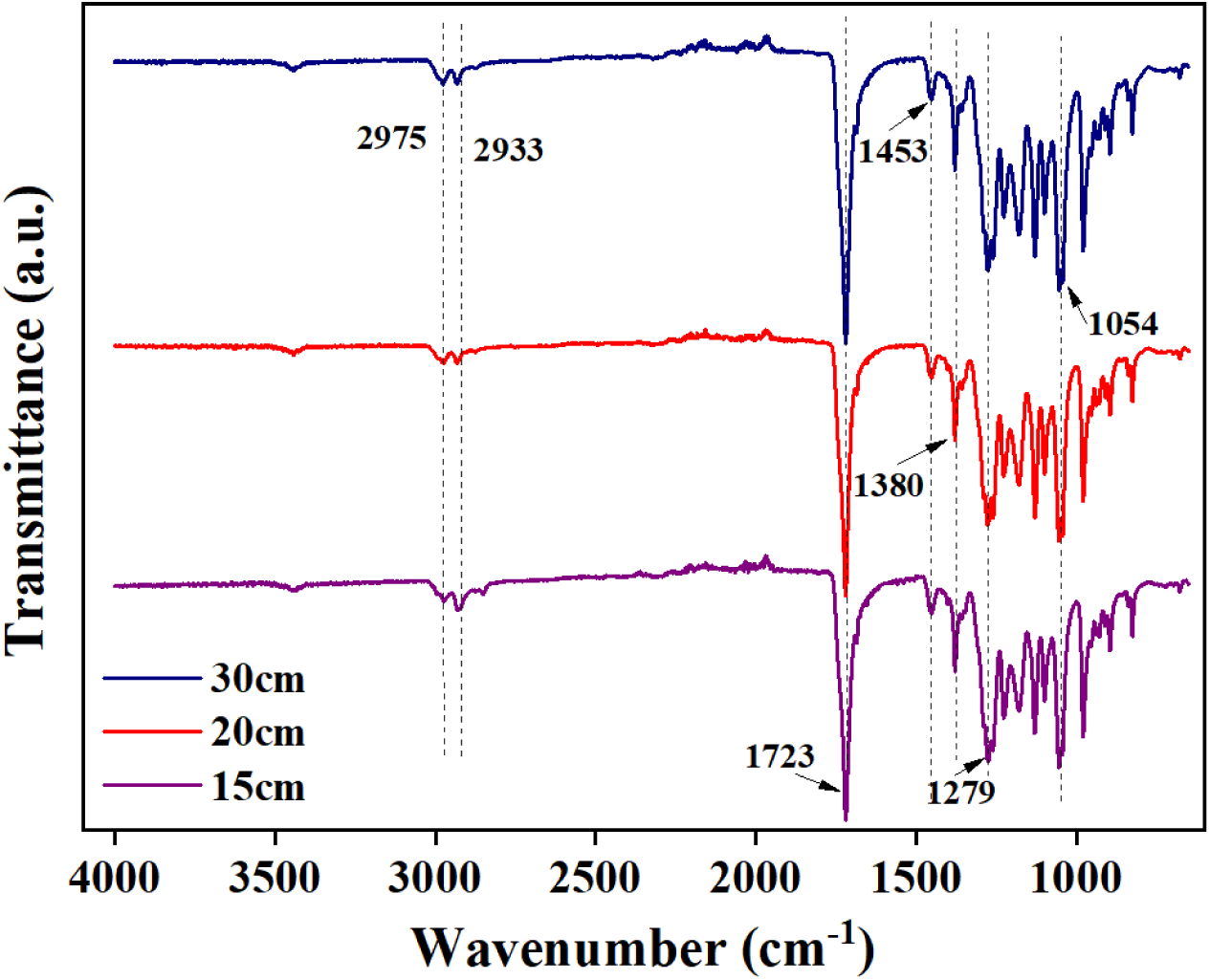
FTIR spectra of the PHBV (B8_CD2) fibers obtained at 15, 20, and 30 cm.

### 3.3 Piezoelectric properties of PHBV fibers

#### 3.3.1 Piezoelectric coefficient of PHBV fibers using the Berlincourt method

After optimising and obtaining the electrospun PHBV fibers, the sample B8_CD2 was selected for further evaluation of their electromechanical coupling through the characterisation of their piezoelectric coefficient (*d*_33_). This evaluation involved the mechanical stimulation of the aligned fibers deposited parallel on aluminum (Al) foil by applying a dynamic force – with an amplitude and a frequency of 0.45 N and 110 Hz, respectively – coupled with a static force of 1N, and measuring the charge generated by the direct piezoelectric effect. It was observed that the *d*_33_ was highly dependent on the morphology of the obtained fibers, as it is shown in **Figure 6a**. In the case of the fibers at 15 cm, the average *d*_33_ was calculated to be 0.117 ± 0.001 pC/N. For the second sample at 20 cm, it was noticed that the *d*_33_ was significantly improved compared to the fibers obtained at 15 cm, with the value to be 25 times higher (2.9 ± 0.2 pC/N) than the *d*_33_ of the fibers obtained at 15cm. This improvement can be attributed to the porous structure of the fibers at 20 cm. The introduction of the air gaps within the fibers structure can enhance further the ability of the material to store more electrical energy.^38^ The fibers obtained at 30 cm demonstrated a *d*_33_ slightly higher than the ones electrospun at 15cm but considerably lower than the fibers collected at 20cm, with a value of 0.16 ± 0.06 pC/N. Even though the fibers at 30cm have porosity on their structure, it is significantly lower than that of the B8_CD2_20 cm and the diameter of the fibers is not uniform and with beads, as indicated in **Figure 4e and 4f**. The combination of pores and bead defects can lead to mechanical instability in the fibrous structure which in turn can reduce the efficiency of piezoelectric excitation. In the next section, we will describe a finite element model (FEM) simulation conducted here to further clarify the effects of the pores on the piezoelectric response of the fibers (**Figure 6b and 6c**).

**Figure 6.**
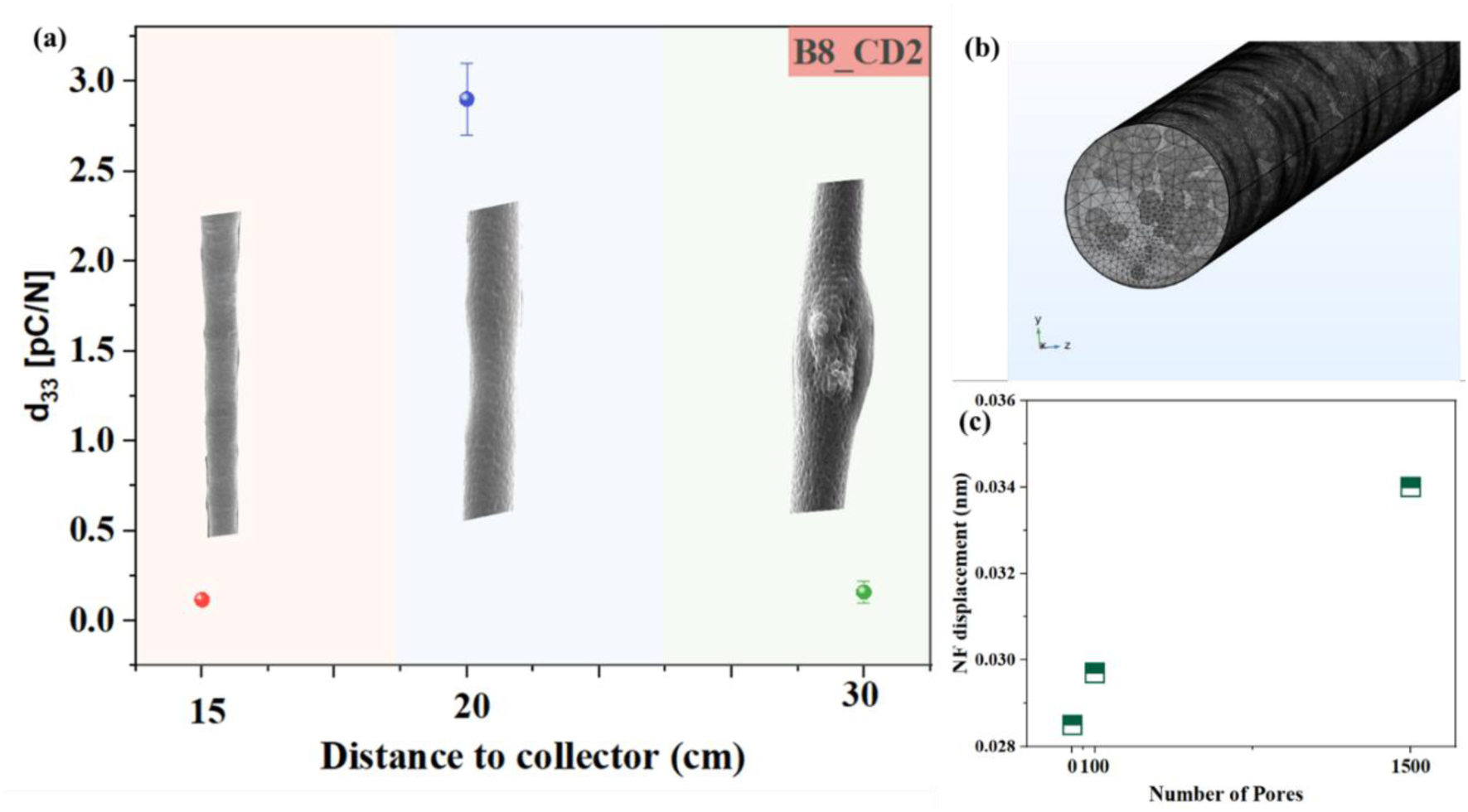
Piezoelectric Response on PHBV Nanofibers with different pores size (a) Piezoelectric coefficient of the B8_CD2 (15cm) electrospun fibrous layers under different distances of the tip-to-collector with inset SEM images of the produced fibers, (b) Image of the 3D simulated fiber with a free tetrahedral mesh and (c) nanofiber displacement distribution as a function of the pores number inside the fiber.

#### 3.3.2 FEM Analysis of Pore Size Effects on Piezoelectric Response in PHBV Nanofibers

A FEM simulation was also carried out to analyze the stress distribution along the y-axis of the fiber, as illustrated in **Video S1**. In these simulations, we have considered that the piezoelectric response of the material is governed by the piezoelectric constitutive equations, which describe the coupling between mechanical and electrical properties. These equations can be expressed in two forms, the stress-charge form (**Equation 4 a and 4b**) and the strain-charge form (**Equation 4 c and 4d**).

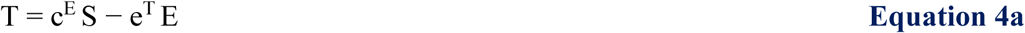

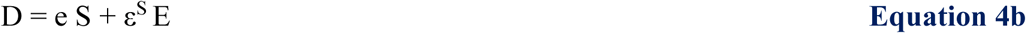

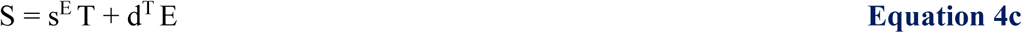

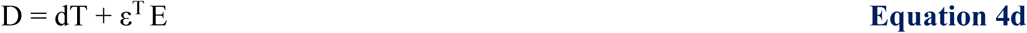

where *T* is the stress tensor, *S* is the strain tensor, *E* is the electric field, and *D* is the electric displacement field. The material properties include the elastic stiffness tensor (c^E^), the piezoelectric coupling matrix (e), and the permittivity matrix (ε^S^) at constant strain. s^E^ represents the elastic compliance matrix at a constant electric field, *d* is the piezoelectric coefficient matrix, and ε^T^ is the permittivity at constant stress.

The fiber was modelled with varying numbers of randomly distributed pores, with diameters ranging from 25 nm to 250 nm, as shown in **Figure 6b**. The resulting displacement of the nanofibrous (NF) structure as a function of pore number is presented in **Figure 6c**, showing a clear increase of the NF displacement with the number of pores. The simulation results indicate that porosity has a significant impact on the piezoelectric properties of the material, with the pore quantity playing a crucial role. Piezoelectric enhancement through porosity is highly sensitive to structural changes. For instance, poor porosity does not provide noticeable improvement, whereas excessive porosity compromises the mechanical integrity of the fibrous structures which in turn weaken their piezoelectric response. This result is in good agreement with previous reports on porous structures, where porosity was concluded to be the main mechanism leading to a higher piezoelectric response of the material.^39^

#### 3.3.3 VNA result and resonance frequency identification

To evaluate the piezoelectric properties of the PHBV fibers, a vector network analyzer was used to measure the frequency response of the fiber with electrodes structure. The impedance (*Z*) and phase spectra were obtained over a wide frequency range to identify resonance characteristics. The measured impedance spectrum, as shown in **Figure 3d**, demonstrates the frequency-dependent electrical response of the PHBV fiber system with copper tape electrodes. The spectrum exhibits distinct features corresponding to the system’s fundamental resonant frequency, which is key indicator of its electromechanical coupling behavior. Analyzing the peak, e.g. the first peak (**Figure 2e**), two primary resonant frequencies are identified, including: series resonance frequency (𝑓_𝑠_) or also known as resonant frequency at 1.76041 ± 0.0002 GHz, and the parallel resonance frequency (𝑓_𝑝_) or also known as anti-resonance at 1.77398 ± 0.0002 GHz. Using **Equation 1**, a coupling coefficient *k*_t_ of 0.1368606 was calculated (i.e., 13.69%)

#### 3.3.4 FEM of pore size distribution on nanofiber response

The eigenfrequency at 1.68 GHz for a single nanofiber of a length and radius of 0.12 cm and 790 nm was derived using an ARPACK solver. The stress distribution across the fiber cross-section and along the fiber are shown in **Figure 7a and 7b**. The primary mode is transverse vibrational as reinforced by the uniform stress distribution along the long axis of the nanofiber.

**Figure 7.**
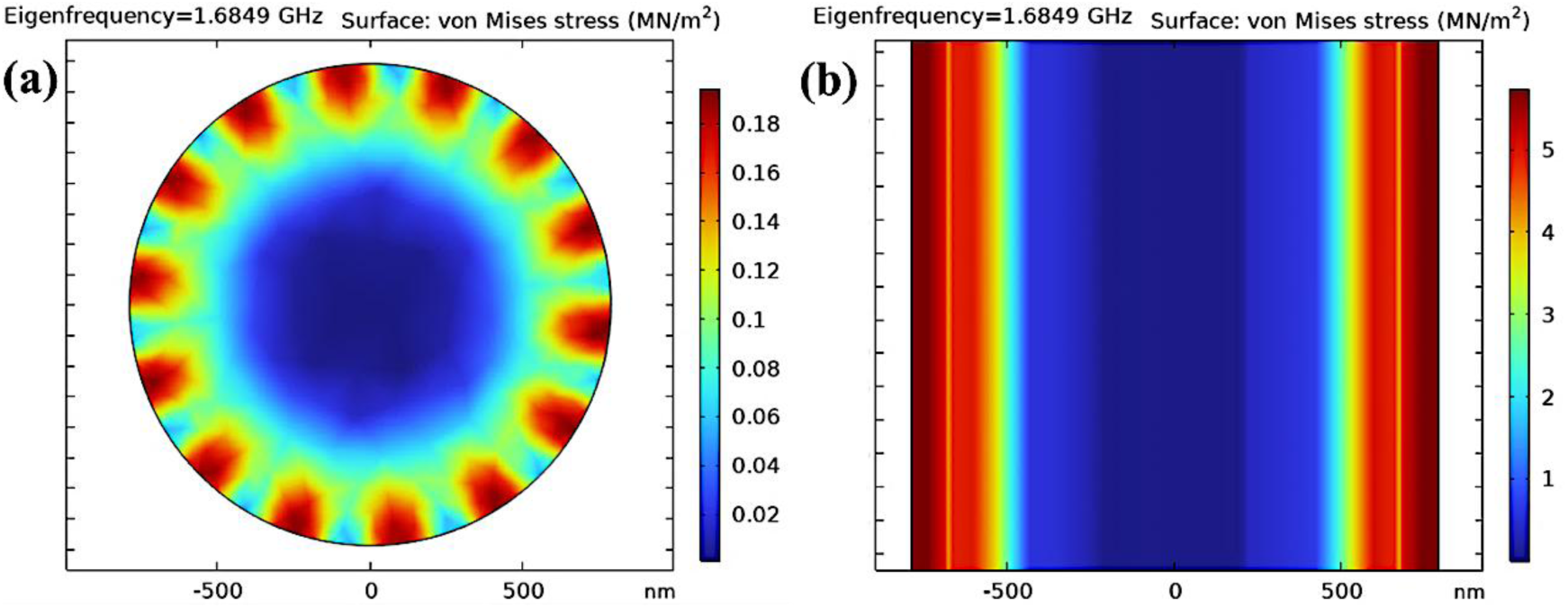
Cross-sectional simulation of the Eigen mode of a single nanofiber (a) perpendicular and (b) parallel to the fiber.

The results obtained from impedance analysis and subsequent calculations confirm the piezoelectric activity of the PHBV fibers. The presence of well-defined resonance peaks, along with a measurable electromechanical coupling factor, demonstrates that the fiber-electrode system effectively converts mechanical stress into electrical signals. The computed *d*_33_ value provides critical insights into the fiber’s piezoelectric efficiency, highlighting its potential for integration into bioelectronics. Nanofibers with a measurable electromechanical coefficient can function as self-powered mechanical sensors or vibration harvesters. By quantifying *d*_33_, you can assess how much electric voltage or current a PHBV-based nanogenerator would produce under mechanical stress or bending. This becomes especially useful in applications like wearable health monitors, structural health sensors, or biodegradable environmental sensors, where sustainable and flexible devices are desired.^40^

### 3.4 Mechanical Properties

The pores have an important role in fiber’s mechanical properties, therefore, the tensile stress-strain curves for the smooth and porous fibers are shown in **Figure S6**. As can be determined from the curves, the difference between the smooth (15 cm) and the porous (20 cm) fibers are distinct through the Young’s modulus (E) for each case. The smooth fibers exhibited a higher E compared to the porous fibers, with calculated values of around 23 MPa and 21 MPa, respectively. These findings align well with the existing literature on PHBV electrospun fibers.^41^ The presence of porosity around the surface of the fibers influences their mechanical behaviour. More specifically, the presence of porous enhances the softness of the fibrous mat, making it more elastic compared to the smooth fibers. Also, at lower stresses, a nonlinear deformation has been observed following by a linear one when the tensile stress increased until reaching the ultimate strength and elongation at break for each case.

Tensile stress-strain measurements were performed for samples of 15, 20, and 30 cm that were rolled into 3D cylinders in dry and wet conditions prior to their evaluation for muscle regeneration to check their capacity to stretch as native muscle in native environments. It is known that muscle undergo strains from 5% for lower intensity training and up to 20% for high intensity training.

In **Figure 8a and b**, 15 and 20 cm fibers undergo higher strain in dry than wet conditions with a maximum strain over 20% strain. We have observed that the fibers after electrospinning in a thin sheet before rolling them in 3D cylinders are withstanding less strain than the one rolled. The enhanced strain tolerance of PHBV fibers rolled into 3D cylinders, compared to thin sheets, can be attributed to improved fiber alignment, structural reinforcement, and more efficient stress distribution.

**Figure 8.**
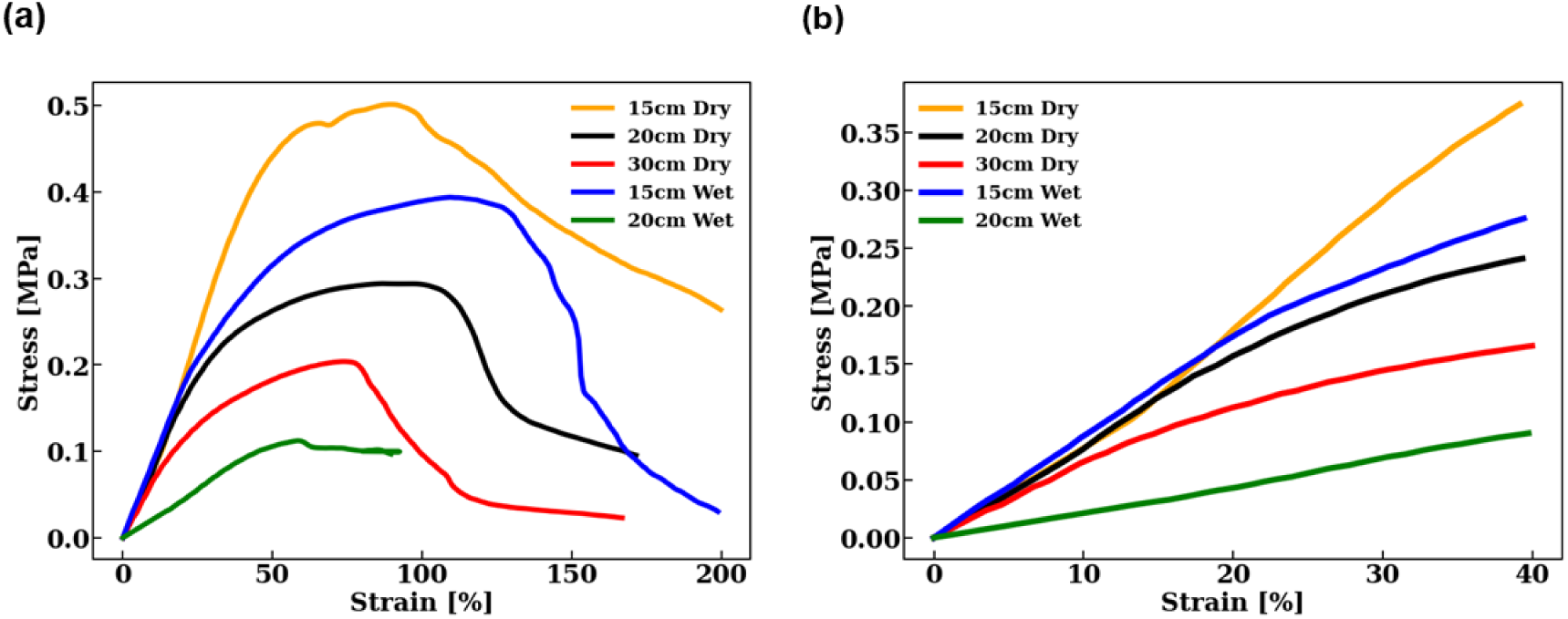
Stress and strain curves from dry (15, 20 and 30 cm samples) and wet (15 and 20 cm samples) tensile testing, respectively. Each curve represents an average of 3 samples, mean ± SD, n=3 (a) The graph showing whole range measurement of stress and strain and (b) The graph depicts the linear region of the whole range stress and strain curves.

### 3.5 Scaffold biocompatibility

Evaluating the biocompatibility of engineered scaffolds represents a critical step in establishing their suitability for tissue regeneration applications. To assess the cytocompatibility of the fabricated electrospun PHBV scaffolds, a live/dead cell viability assay was conducted at three distinct time points: 1, 3, and 7 days. Two different scaffolds obtained by variations in the electrospinning tip-to-collector distance (15 cm and 20 cm), were evaluated, both with and without laminin (LM) incorporation.

Quantitative analysis of cell viability revealed consistently high levels of viable cells across all tested conditions (**Figure 9**). This finding strongly indicates the excellent biocompatibility of the PHBV scaffolds. Notably, scaffolds fabricated at a 15 cm tip-to-collector distance exhibited superior cell viability, consistently exceeding 80% at each time point. This suggests that these scaffolds provide an optimal microenvironment for cell attachment, survival, and proliferation.

**Figure 9.**
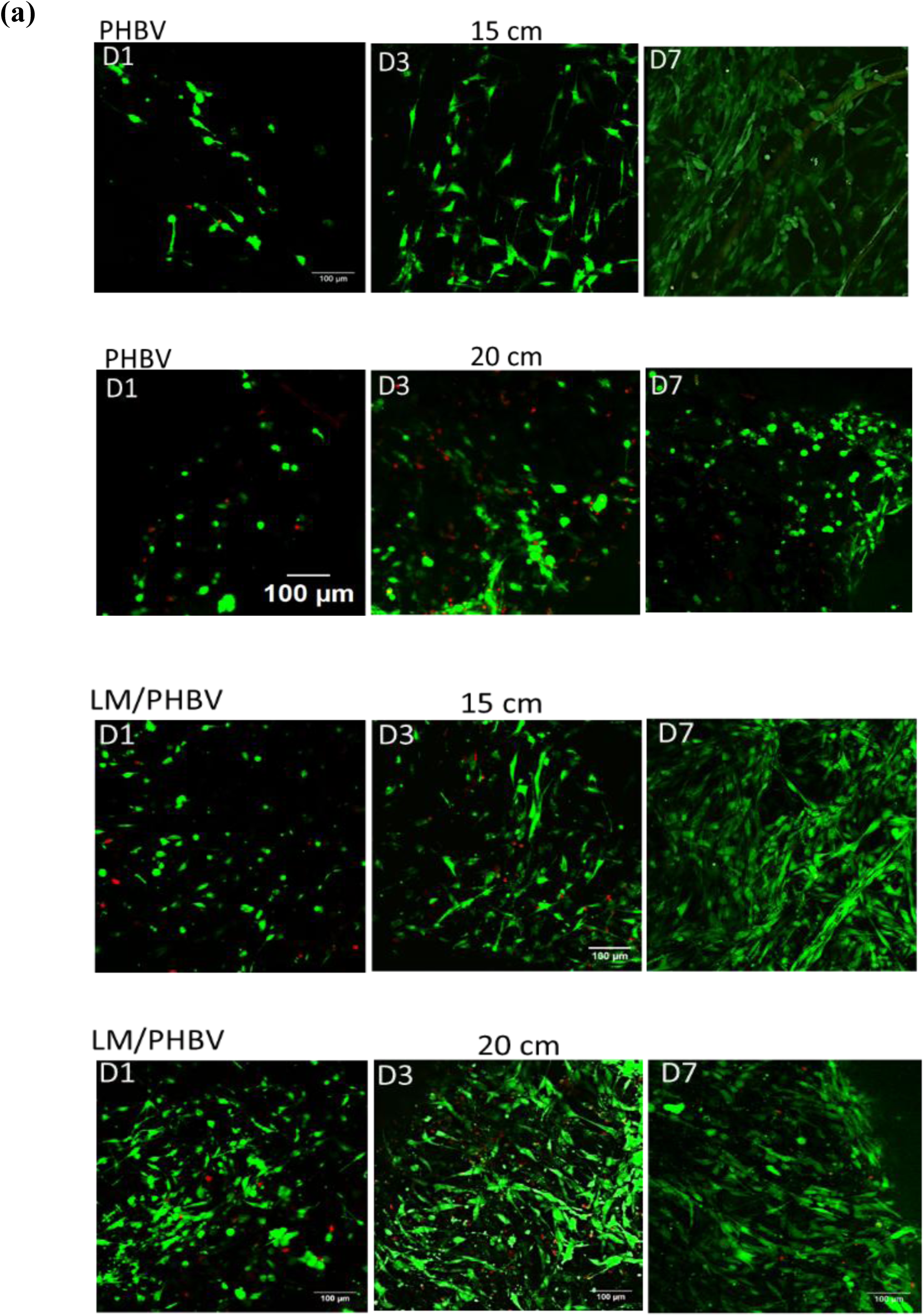
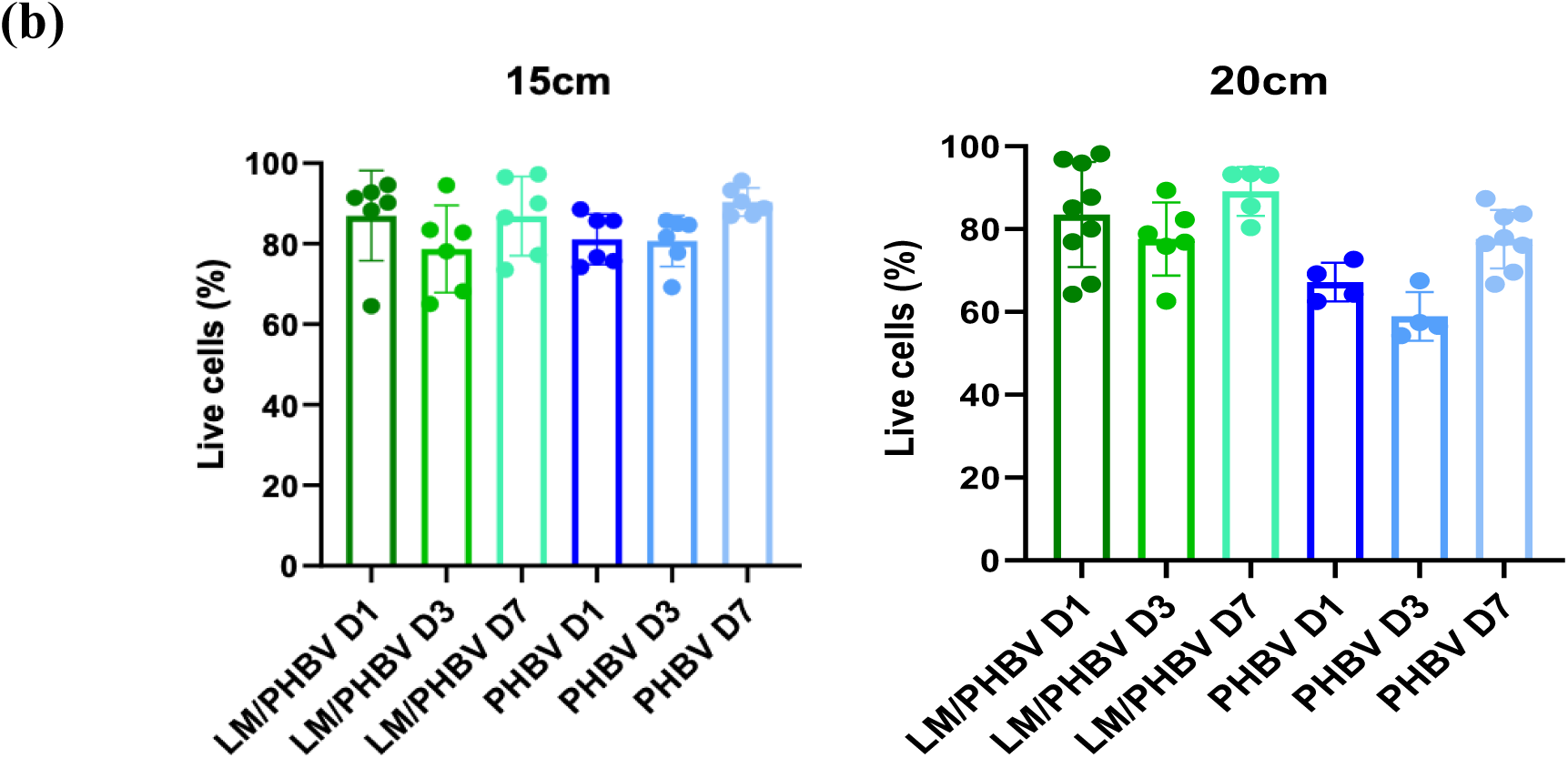
Viability of C2C12 cells on PHBV scaffolds with and without laminin treatment for 1, 3, and 7 days for 15 and 20 cm. (a) Representative fluorescence confocal images of live/dead staining (live cells: green, dead cells: red, scale bar: 100 μm, n = 3) and (b) Quantification of live cells (%) from the images of 15 and 20 cm fibers. Statistics are shown as mean ± standard deviation. Data were analyzed by one-way ANOVA (α = 0.05). ****p < 0.0001.

Interestingly, a slight decrease in cell viability was observed on scaffolds fabricated at a 20 cm tip-to-collector distance (**Figure 9b**). Confocal images (**Figure 9a**) revealed an altered fibers surface in these scaffolds, characterised by less uniform cell distribution and increased cell clumping compared to the 15 cm scaffolds. This altered morphology could potentially hinder cell-cell interactions and impact overall cell behaviour, contributing to the observed difference in cell viability. These results highlight the critical role of scaffold architecture in dictating not only cell viability but also the spatial distribution of cells within the scaffold.

### 3.6 Cell-material interaction

To evaluate the cell-material interaction, vinculin expressions were assessed after C2C12 was seeded onto the scaffold for 3h. In **Figure 10a**, it has been observed that C2C12 cells seeded on PHBV scaffolds have smaller and rounded morphology with weaker focal adhesion expression, whereas the cells on LM/PHBV scaffolds were found to have elongated morphology with stronger adhesion and long wedge shape focal adhesion which demonstrated the need of the extracellular matrix protein for the cells to adhere. It has been also confirmed that cells on LM/PHBV have a higher aspect ratio on 15 cm compared to 20 cm with cells following similar trend when laminin is not present demonstrating that fibers surface is different with cells more elongate adhesions on 15 than 20 cm (see supplementary (graphs aspect ratio) **Figure S7**). More, quantitative analysis (**Figure 10b**) reveals that the total number of detected focal adhesion (FA) per cell between cells seeded on 15 cm and 20 cm PHBV scaffold has no difference while higher expression of FA per cell were observed from cells seeded on LM/PHBV 15 cm compared to 20 cm.

**Figure 10.**
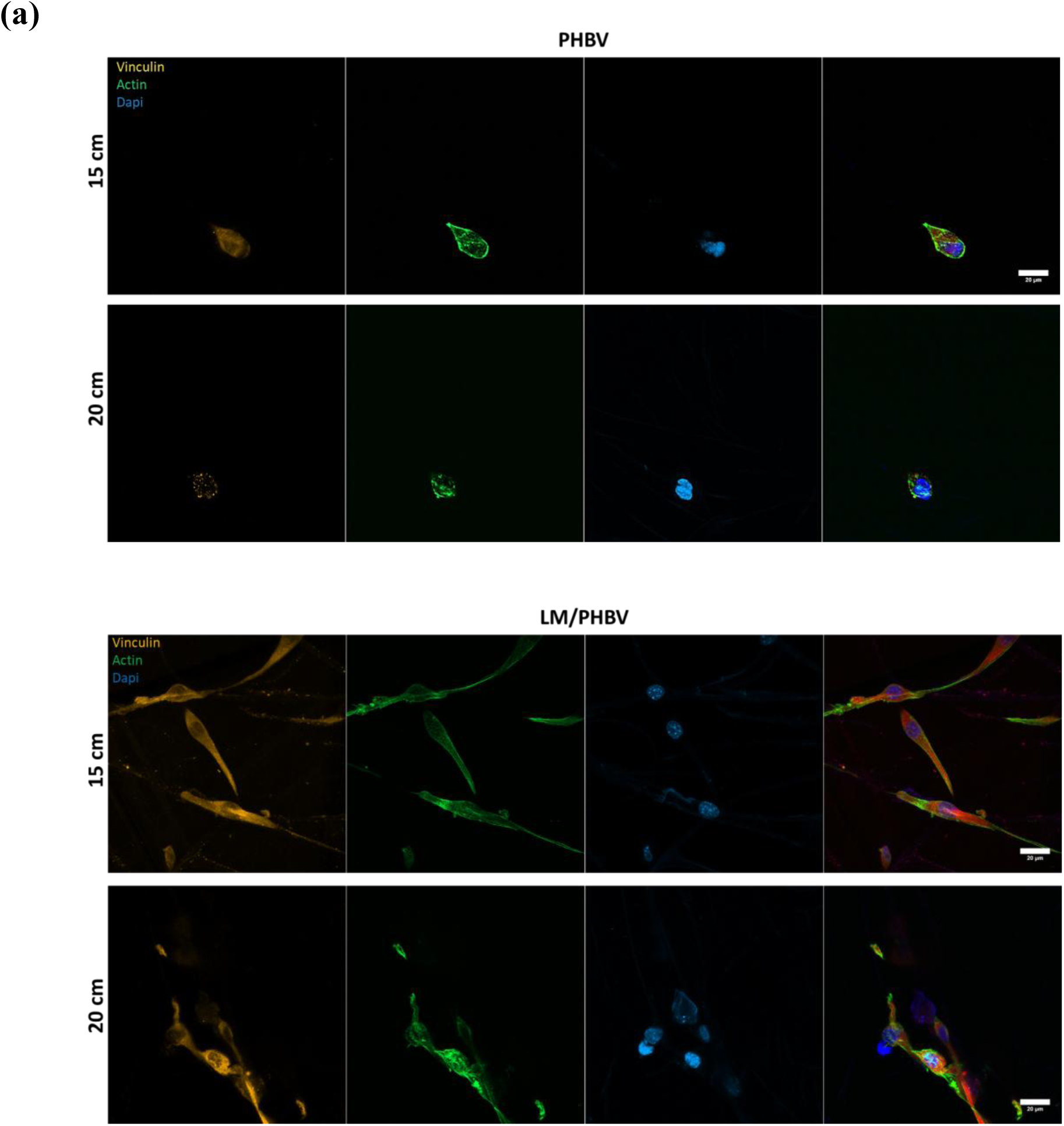
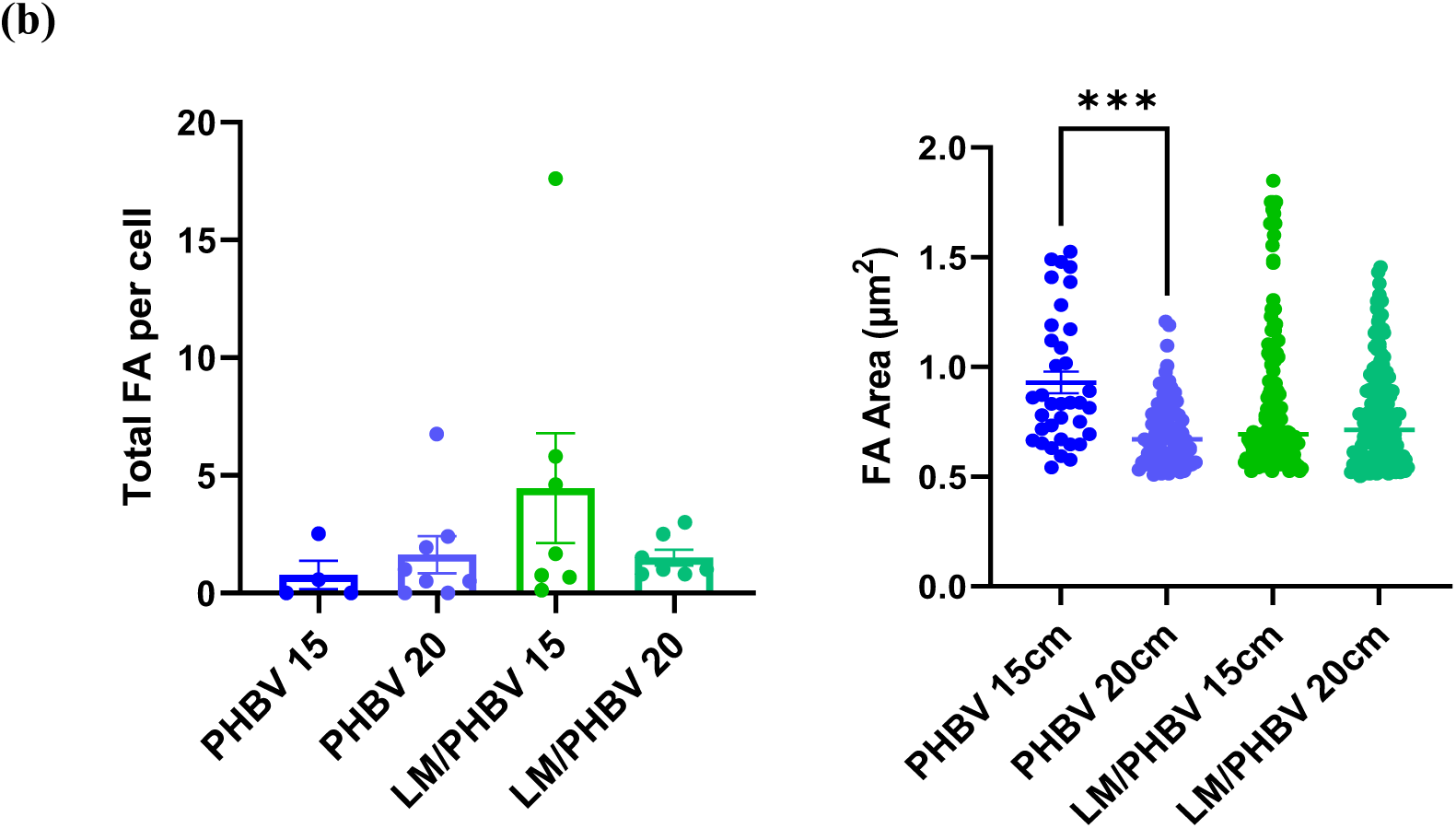
(a) Representative confocal images of C2C12 cells cultured on 15cm and 20cm PHBV and LM/PHBV electrospun scaffolds. Cells were fixed and stained with phalloidin-488 for actin cytoskeleton (green), anti-vinculin for FAs (orange/gold), DAPI for Nucleus (Blue). Scale bar: 20 µm. (b) Quantification of total FA per cell and FA area in C2C12 cells cultured on 15cm and 20cm PHBV and LM/PHBV scaffolds. Each data point corresponds to observation from SEM (n < 4 for each condition).

### 3.7 Muscle regeneration

The potential of PHBV-based scaffolds to support and promote myogenic differentiation was investigated using C2C12 cells cultured on two distinct scaffold variants (referred to as ’15cm’ and ’20cm’) at two time points: day 5 and day 10. Myotube formation, a critical early indicator of muscle regeneration, was assessed via myosin differentiation staining.

At day 5, immunofluorescence imaging (**Figure 11a**) and subsequent quantification (**Figure 11b**) revealed significant myotube formation on both LM/PHBV and PHBV scaffolds. Specifically, 15cm scaffolds demonstrated approximately 40% myotube formation when cultured in induction media. While both LM/PHBV and PHBV scaffolds displayed comparatively lower myotube formation at day 5 relative to a positive control, they still promoted considerable differentiation, with over 5% of total cells differentiating into myotubes after only 5 days of culture. This early differentiation indicates the intrinsic potential of these scaffolds to support myogenic differentiation.

**Figure 11.**
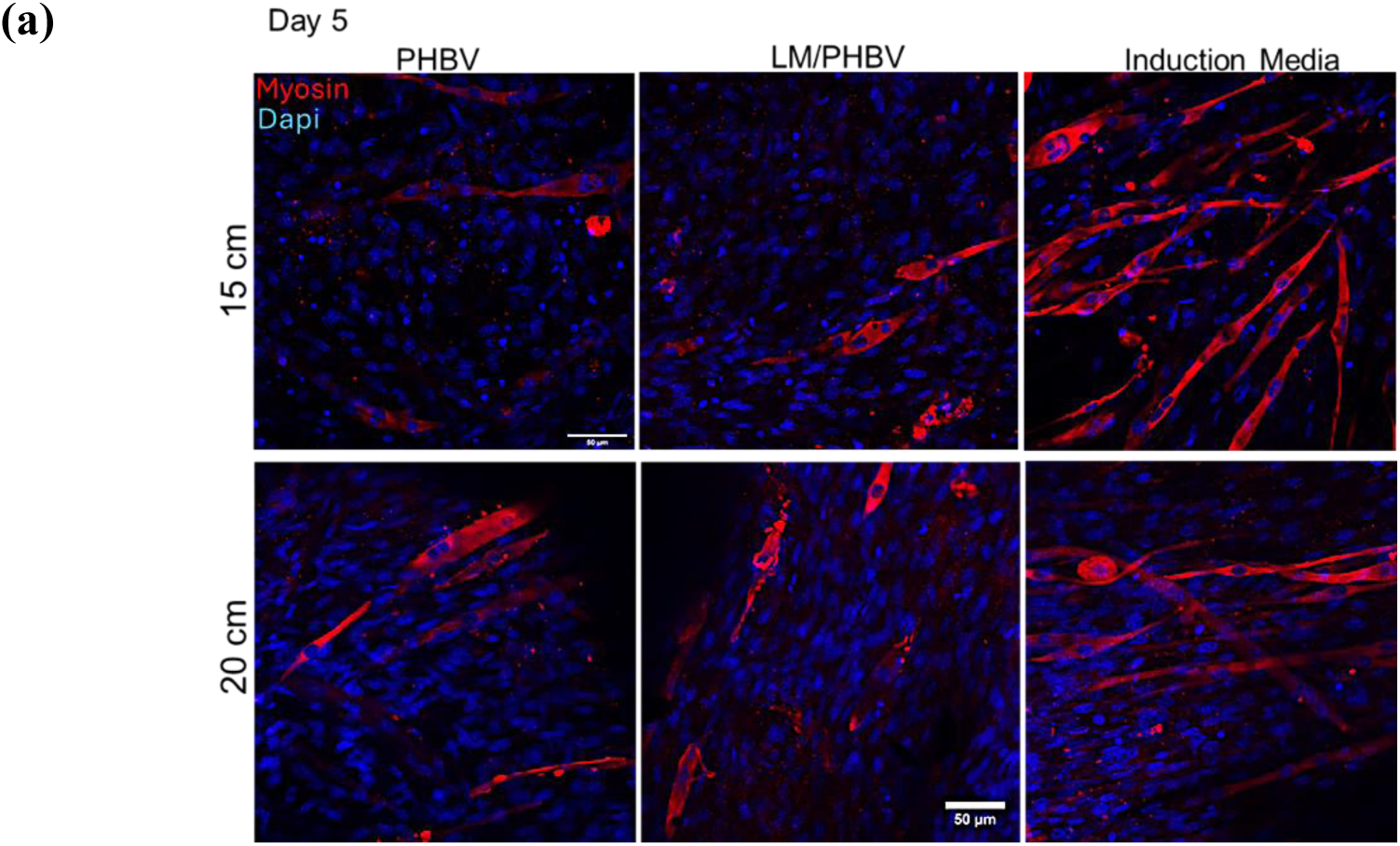
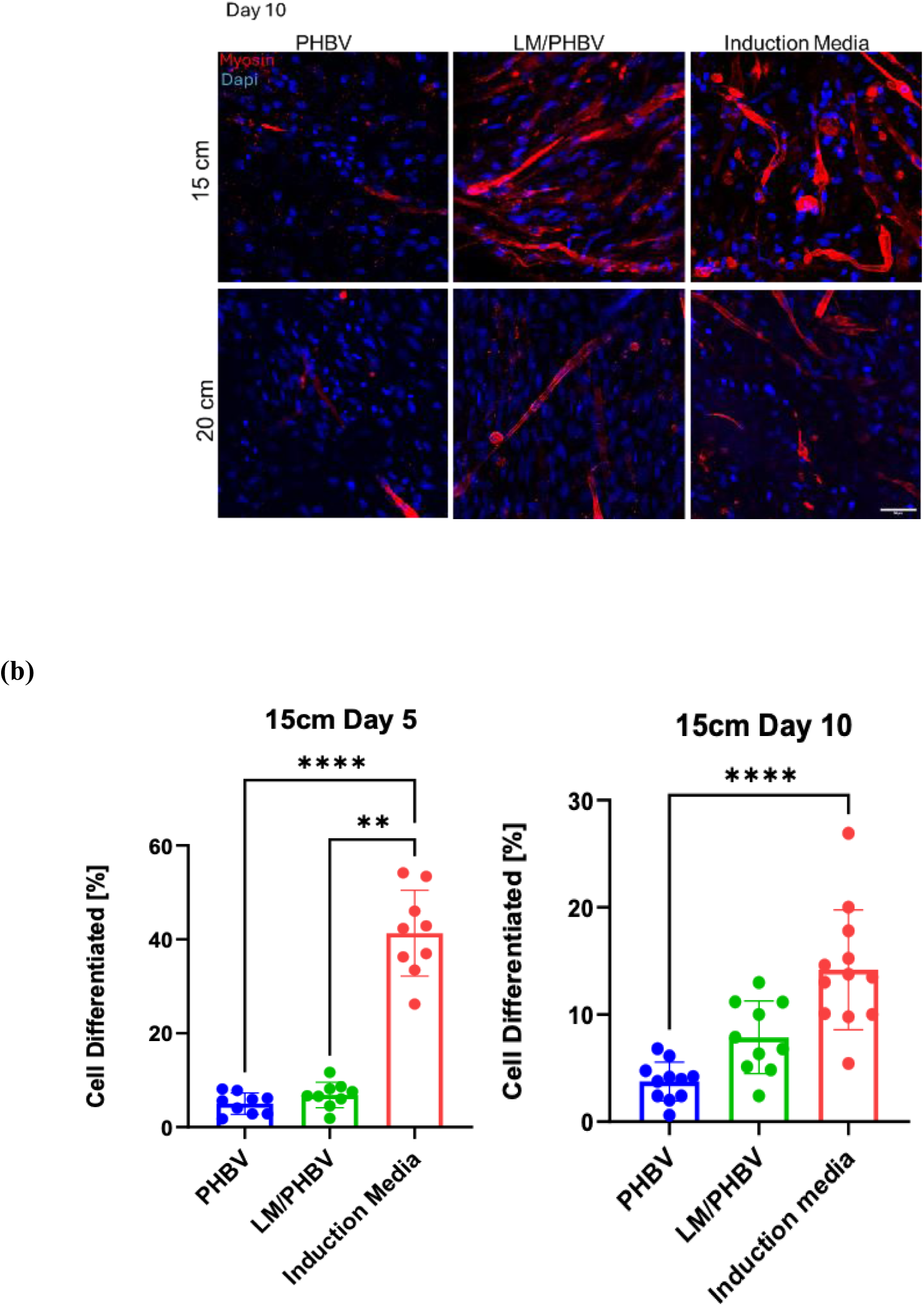
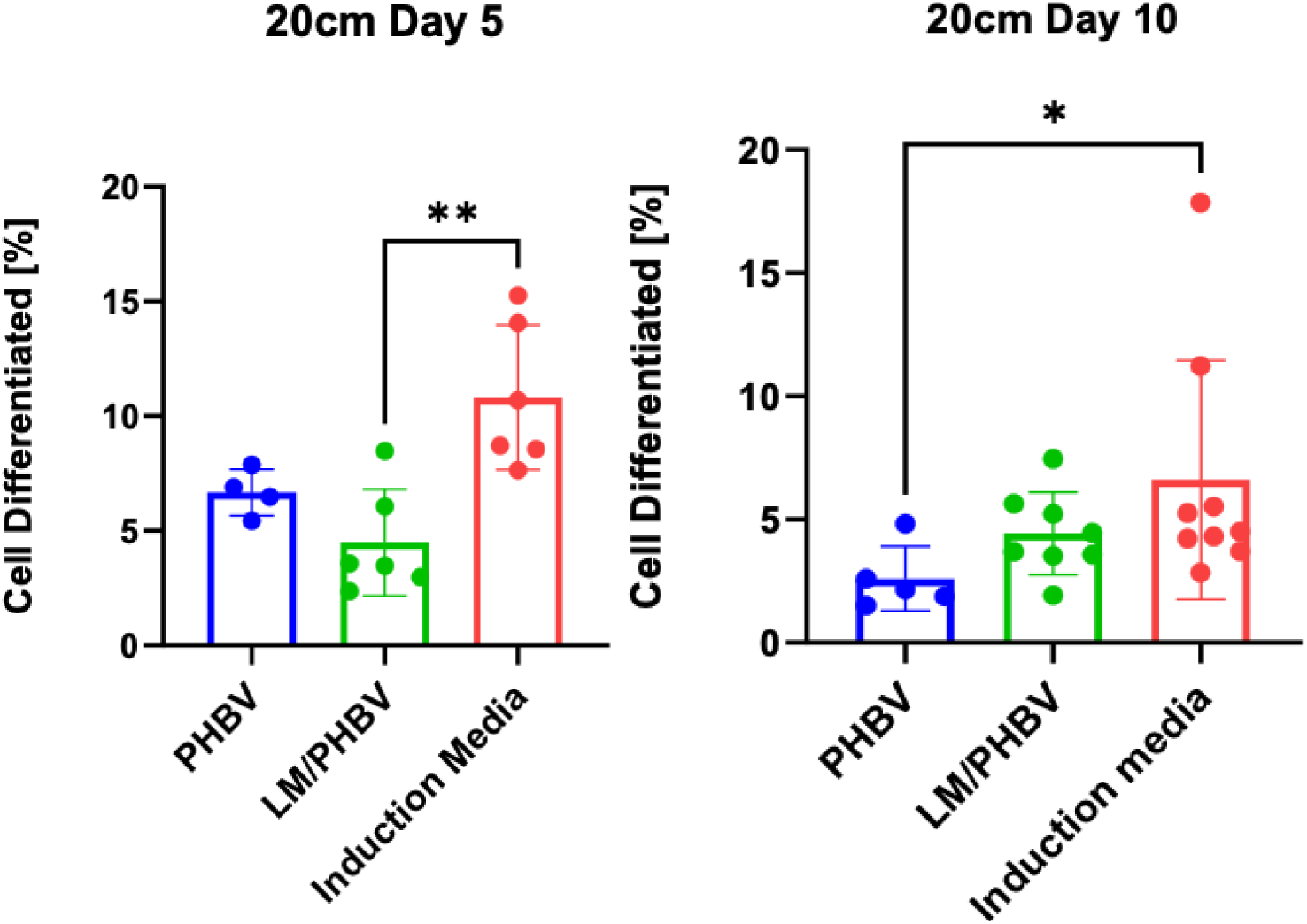
LM/PHVB scaffolds promotes myogenic differentiation in vitro. (a) Immunofluorescence images for evaluation of myotube formation using myosin 4 antibody (MF-20) as specific muscle marker (purple), and counterstained with DAPI (blue) for nuclei visualization and (b) image analysis quantification of total cell number and the percentage of myotube fusion. Note that myotubes were defined and counted as cells with three or more aligned nuclei. Scale bar for confocal images (left column) 50 μm, n = 5 images from 3 different biological replicas per condition. Statistics are shown as mean ± standard deviation. Data was analyzed by ANOVA (α = 0.05). ****p < 0.0001, **p < 0.01, and *p < 0.5.

Analysis of the scaffold 20cm at day 5 also contributed to the overall assessment of scaffold performance (**Figure 11a**). Interestingly, for 15cm scaffolds cultured in induction media, myotube formation at day 10 was notably lower, showing only around 10% cell differentiation, compared to its performance on day 5. This observed decrease in differentiation over time could potentially be attributed to specific characteristics of the scaffold fibers, such as porosity, which might modulate or slow the progression of myogenic differentiation beyond initial induction. Furthermore, by day 10, 15cm scaffolds demonstrated a better capacity to promote myogenic differentiation compared to 20cm scaffolds when both were assessed relative to positive control, suggesting an increased potential over longer culture periods.

### 3.8 LMPHBV scaffold promoting myogenic differentiation in-vitro

The myogenic potential of C2C12 cells cultured on PHBV and LM/PHBV was further evaluated for the expression of specific myogenic differentiation markers such as myogenin and Myo D genes representing early and late markers. C2C12 cells were seeded onto LM/PHBV scaffolds for 5 days and expression of myogenin and MyoD genes specific for myogenic differentiation were evaluated. In **Figure 12a**, we have observed an upregulation of MyoD and myogenin, when compared with positive control.

**Figure 12.**
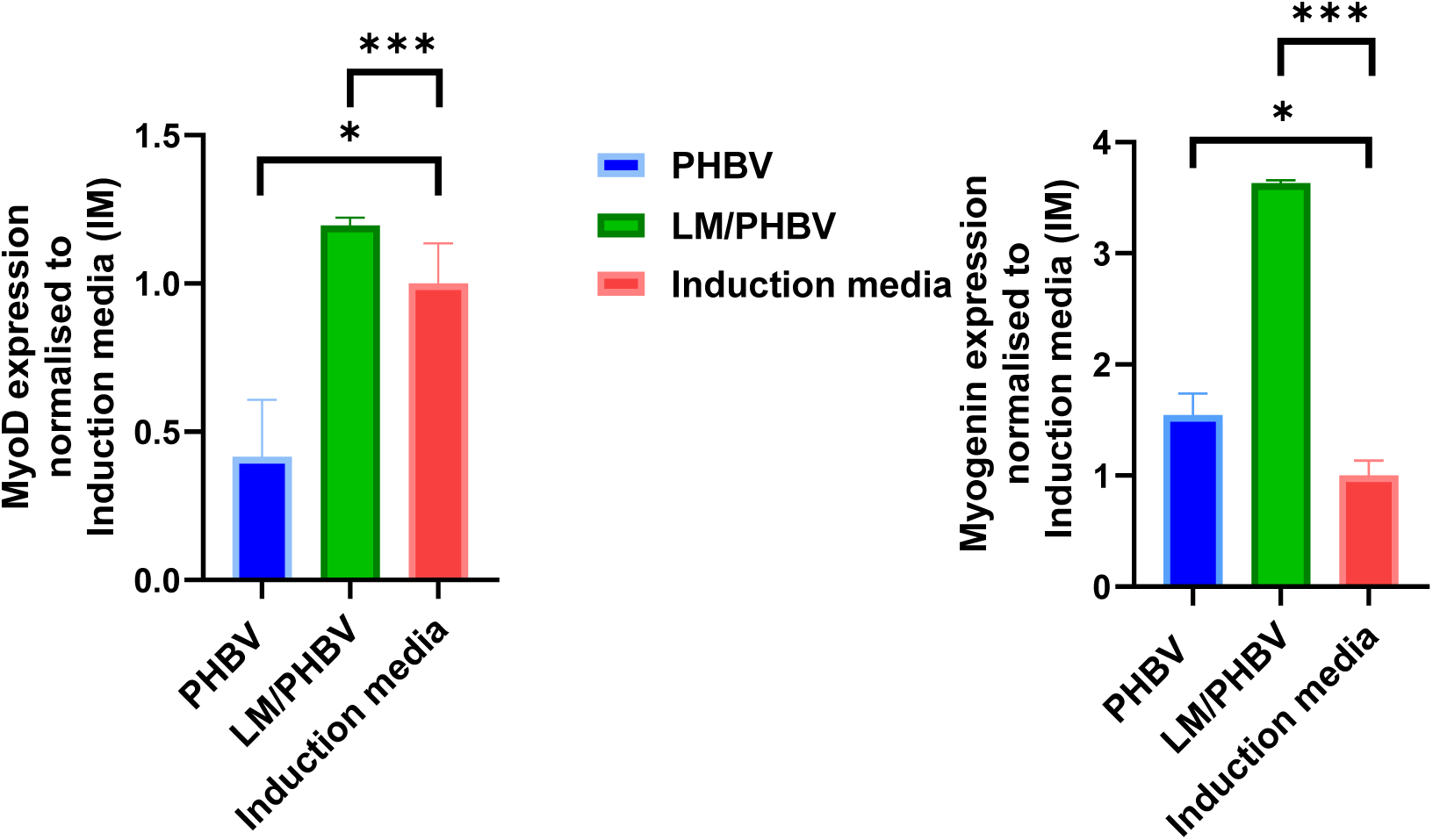
LM/PHBV scaffolds promoted myogenic differentiation. Real-time qPCR analysis for markers of myogenesis (MyoD and Myogenin) after 4 days. Each condition was normalized to the myogenic media (mean ± SD, n = 3, one-way ANOVA with Tukey multiple comparison), where ***p = 0.001, *p = 0.5 showed an upregulation compared with positive control.

Overall, as depicted in **Figure 12b**, LM/PHBV scaffolds have been shown to promote myogenic differentiation in vitro at both protein and gene expression levels in static conditions. We hypothesize that mechanically/electrically stimulated LM/PHBV scaffolds will enhance or accelerate the myogenic regeneration due to the known importance of the muscle cells to contract and stretch.

## 4. Conclusion

This study establishes electrospun PHBV fibrous scaffolds as promising biomaterials for regenerative medicine, with a particular focus on muscle tissue engineering. Through systematic optimization of processing parameters, we successfully fabricated scaffolds with controlled porosity and fiber morphology, significantly influencing their functional properties. Notably, the introduction of porosity resulted in a remarkable 25-fold enhancement of the *d*_33_ compared to non-porous scaffolds. Quasi-static (Berlincourt) and dynamic (VNA) response of the PHBV NFs were derived highlighting physics-based insights into material properties. Furthermore, the simulation framework, exploiting the randomised pore distributions, further confirms the enhancement of piezoelectric response due to the newly introduced stress distributions around the pores within the PHBV nanofibers.

This enhanced piezoelectricity, coupled with suitable mechanical properties mimicking native muscle tissue, suggests a high potential for these scaffolds to promote favorable cellular responses, including cell adhesion, proliferation, and differentiation. While our current focus lies in muscle tissue engineering, the versatility of PHBV, its biocompatibility, and the ability to tailor its properties through electrospinning suggest broad applicability in addressing various tissue regeneration challenges, including bone, cartilage, and nerve repair.^42^ This study lays the groundwork for future investigations into the use of these piezoelectric PHBV scaffolds in a wider range of tissue engineering applications.

## AUTHOR INFORMATION

Corresponding Author

* (O.D.)

## Author Contributions

X. K. and G. B. contributed equally to this work.

## ACKNOWLEDGMENT

SEM imagining was carried out in Electron Microscopy Unit/CAF College Shared Research Facilities, College of Medical, Veterinary & Life Sciences (MVLS) by Margaret Mullin. The authors also thank Prof. Scott Ramsay and Miss Crystal Liu from University of Toronto for providing the tensile tester. Dr. Giuseppe Ciccone for technical assistance and advice with the tensile tester. Dr Manuel Pelayo Garcia for technical assistance on the electrical characterization of the fibers. Dr. Afesomeh Ofiare for assistance in VNA calibration.

## ABBREVIATIONS

Poly(3-hydroxybutyrate-co-3-hydroxyvalerate) (PHBV), Fibrous piezoelectric scaffolds (FPS), Laminin (LM), Extracellular matrix (ECM), focal adhesion (FA),

