## Supplementary Information for "Three-Dimensional Piezoelectric Fibrous Constructs for Muscle Regeneration"

ASSOCIATED CONTENT

### SUPPLEMENTARY

**Table S1:** Experimental details for the preparation of PHBV solutions

| Provider | Concentration (wt.%) | Solvent 1 | Solvent 2 | Volume per volume ratio (v/v) | Acronym |
|---|---|---|---|---|---|
| **Biopol** | 5 | Chloroform | DCM | 1:1 | B5_CD1 |
| | | Chloroform | DCM | 4:1 | B5_CD2 |
| | | DCM | DMF | 9:1 | B5_DF |
| | | Chloroform | DMF | 9:1 | B5_CF |
| | 8 | Chloroform | DCM | 4:1 | B8_CD2 |
| | 10 | Chloroform | DCM | 4:1 | B10_CD2 |
| **Sigma Aldrich** | 5 | Chloroform | DCM | 1:1 | SA5_CD1 |
| | | Chloroform | DCM | 4:1 | SA5_CD2 |

| | | | | | |
|---|---|---|---|---|---|
| | | DCM | DMF | 9:1 | SA5_DF |
| | | Chloroform | DMF | 9:1 | SA5_CF |
| | 8 | DCM | DMF | 9:1 | SA8_DF |
| | 10 | DCM | DMF | 9:1 | SA10_DF |

**Table S2**: Applied voltages during electrospinning for all solutions

| PHBV Fibers | Applied voltage (kV) | PHBV Fibers | Applied voltage (kV) |
|---|---|---|---|
| **B5_CD1** | 13 | **SA5_CD1** | 15 |
| **B5_CD2** | 14 | **SA5_CD2** | 15 |
| **B5_DF** | 13 | **SA5_DF** | 13 |
| **B5_CF** | 15 | **SA5_CF** | 12 |
| **B8_CD2** | 15 | **SA8_DF** | 10 |
| **B10_CD2** | - | **SA10_DF** | 14 |

**Table S3**: Properties used in the FEM simulation for the PHBV fibers

| Material Parameters | Smooth Fibers | Porous Fibers |
|---|---|---|
| **Density [kg/m3]** | 1250 | |
| **Poisson's ratio** | 0.35 | |
| **Young's modulus [MPa]** | 22.68 | 21.03 |
| **d33 [pC/N]** | 0.117 | 2.9 |

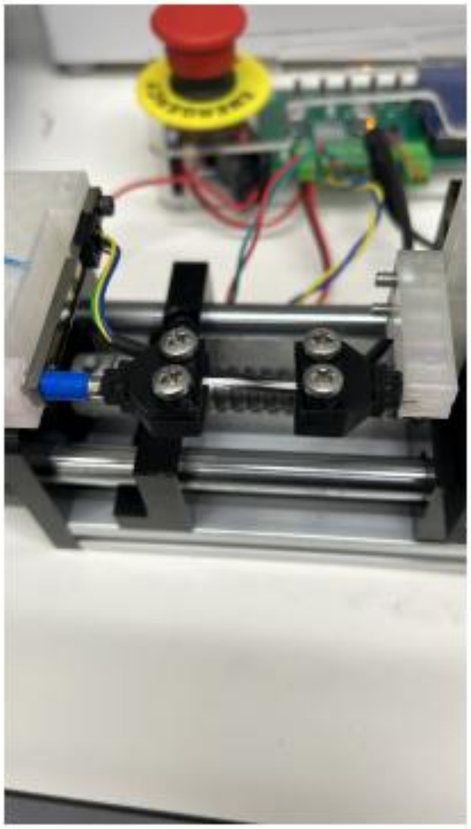

**Figure S1.** Mechanical tensile test setup

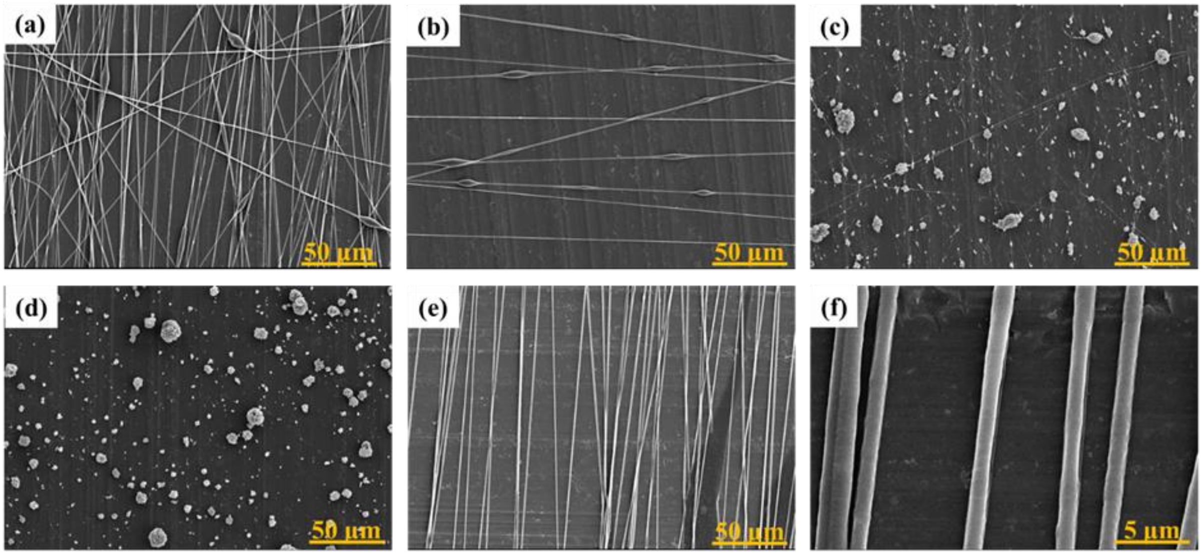


**Figure S2.** SEM images of the PHBV fibers obtained from Biopol (a) B5_CD1, (b) B5_CD2, (c) B5_DF, (d) B5_CF, (e) B8_CD2 and (f) higher magnification of B8_CD2.

**Table S3**: Morphological analysis of electrospun PHBV fibers for all solutions and processing conditions.

| PHBV Fibers | Average diameter of Fibers [µm] | Average diameter of Beads [µm] | Average diameter of Pores [µm] |
|---|---|---|---|
| **B5_CD1** | 0.91 ± 0.31 | 2.82 ± 1.04 | - |
| **B5_CD2** | 0.97 ± 0.21 | 3.39 ± 1.78 | - |
| **B5_DF** | No fibers | 1.65 ± 1.19 | - |
| **B5_CF** | No fibers | 8.05 ± 3.34 | - |
| **B8_CD2_15cm** | 1.27 ± 0.19 | - | - |
| **B8_CD2_20cm** | 1.58 ± 0.44 | - | 0.29 ± 0.15 |
| **B8_CD2_30cm** | 2.09 ± 0.66 | 8.46 ± 4.72 | 0.15 ± 0.05 |
| **SA5_CD1** | No fibers | 11.88 ± 7.65 | - |
| **SA5_CD2** | No fibers | 12.16 ± 7.65 | - |
| **SA5_DF** | 0.44 ± 0.16 | 2.4 ± 1.25 | - |
| **SA5_CF** | No fibers | 5.41 ± 2.21 | - |
| **SA8_DF** | 0.498 ± 0.2 | 2.81 ± 1.11 | - |
| **SA10_DF** | 0.38 ± 0.18 | 4.61 ± 2.17 | - |

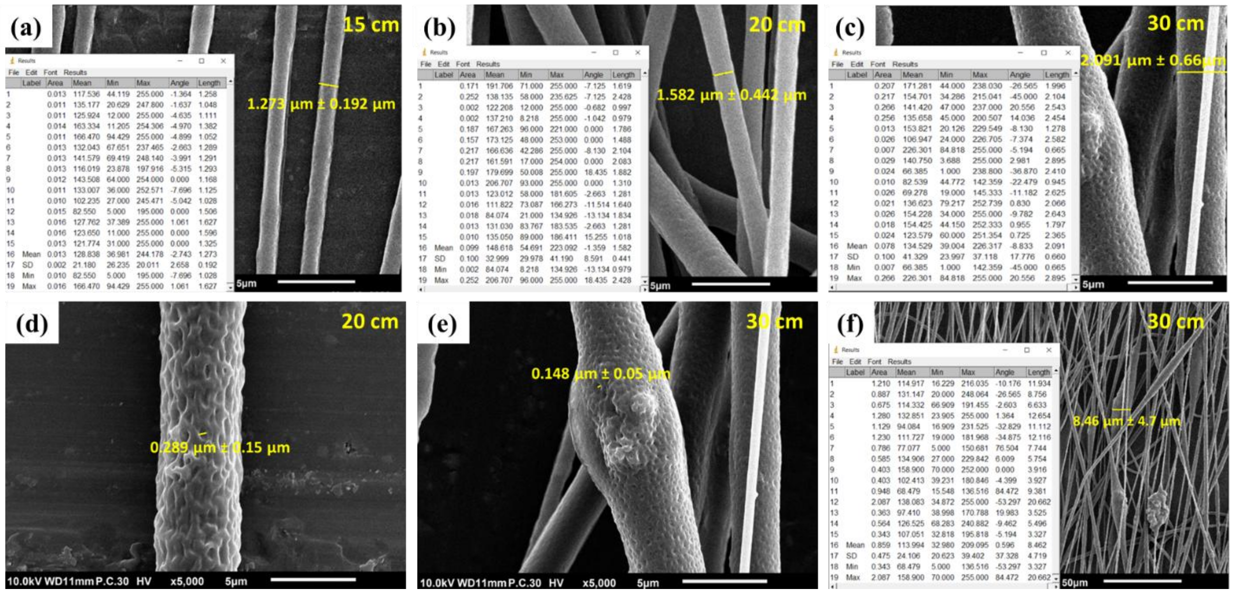

**Figure S3.** Statistical analysis of B8_CD2 fiber diameters at different tip-to-collector distances: (a) 15 cm, (b) 20 cm, and (c) 30 cm. Pore diameter analysis of B8_CD2 fibers at (d) 20 cm and (e) 30 cm. (f) Average bead diameter of B8_CD2 fibers at 30 cm

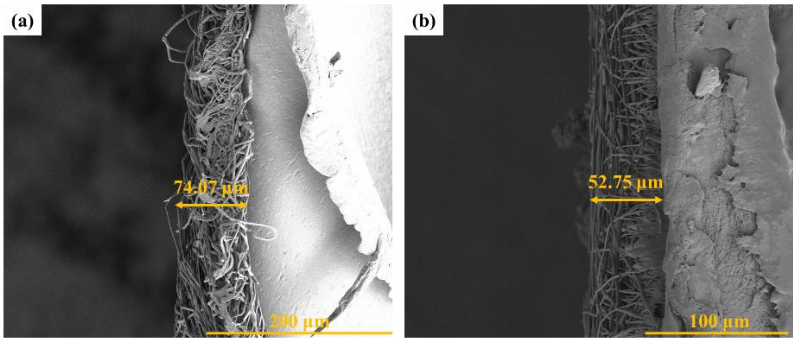


**Figure S4**. Cross-sectional SEM images of the B8_CD2 fibers obtained at (a) 15 cm and (b) 20 cm.

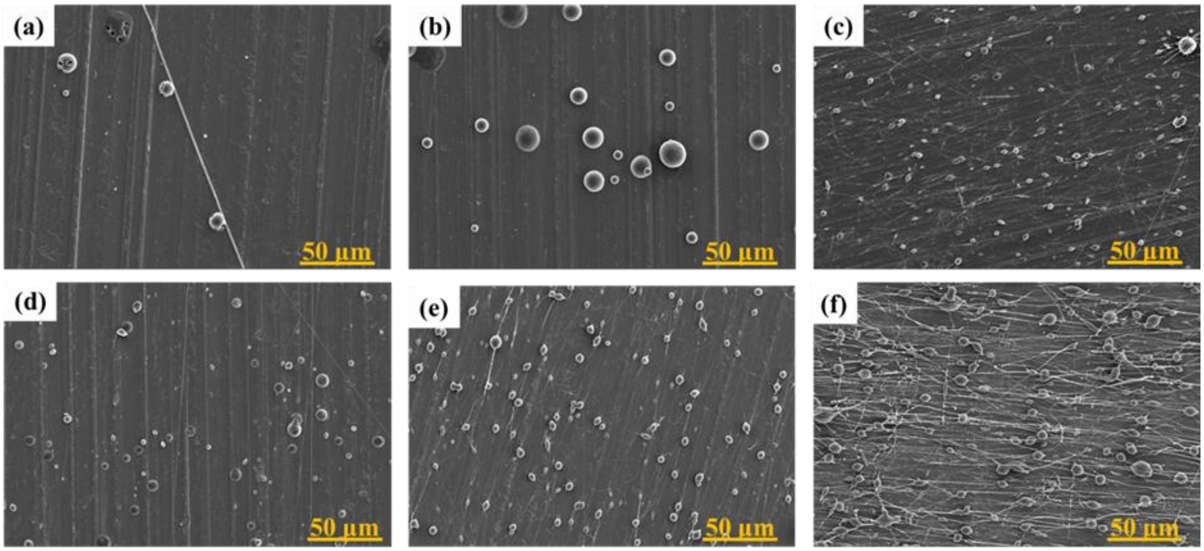


**Figure S5.** SEM images of the PHBV fibers obtained from Sigma Aldrich (a) SA5_CD1, (b) SA5_CD2, (c) SA5_DF, (d) SA5_CF, (e) SA8_DF and (f) SA10_DF.

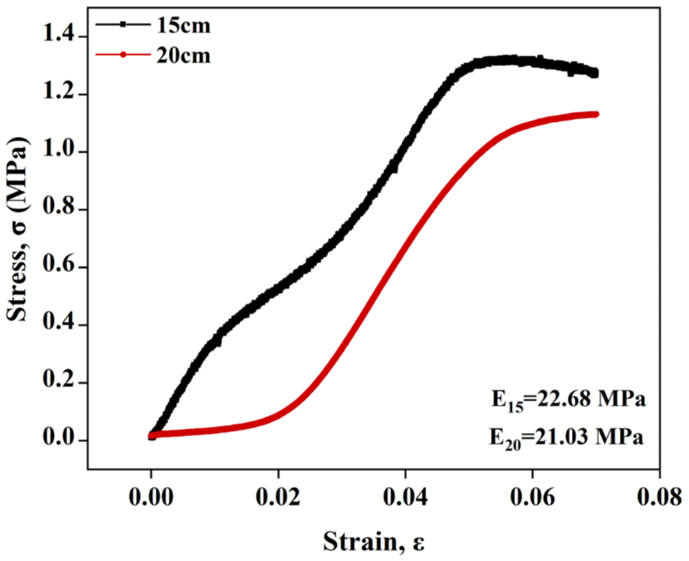


**Figure S6.** Tensile stress-strain curves for the sheet of PHBV fibers electrospun at 15 and 20 cm.

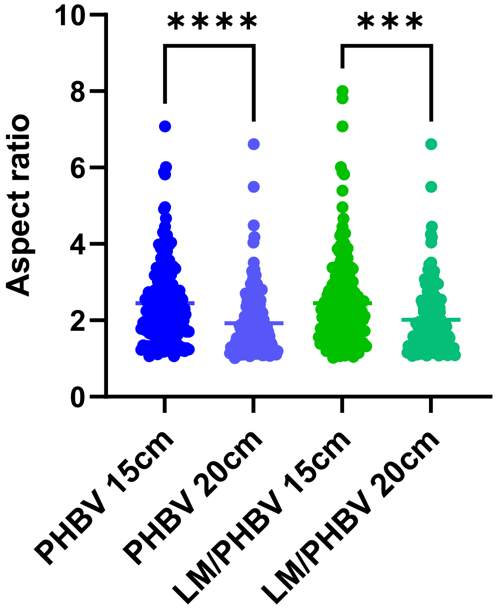


**Figure S7.** Aspect ratio of foal adhesions assessment of C2C12 on PHBV and LM/PHBV with 15 and 20 cm fibers.
